# Social learning strategies for matters of taste

**DOI:** 10.1101/170191

**Authors:** Pantelis P. Analytis, Daniel Barkoczi, Stefan M. Herzog

## Abstract

Most choices people make are about “matters of taste” on which there is no universal, objective truth. Nevertheless, people can learn from the experiences of individuals with similar tastes who have already evaluated the available options—a potential harnessed by recommender systems. We mapped recommender system algorithms to models of human judgment and decision making about “matters of fact” and recast the latter as social learning strategies for “matters of taste.” Using computer simulations on a large-scale, empirical dataset, we studied how people could leverage the experiences of others to make better decisions. Our simulation showed that experienced individuals can benefit from relying mostly on the opinions of seemingly similar people; inexperienced individuals, in contrast, cannot reliably estimate similarity and are better off picking the mainstream option despite differences in taste. Crucially, the level of experience beyond which people should switch to similarity-heavy strategies varies substantially across individuals and depends on (i) how mainstream (or alternative) an individual’s tastes are and (ii) the level of dispersion in taste similarity with the other people in the group.

## 1 Introduction

Where should I go for my next vacation? Which novel should I read next? Most choices people make are about “matters of taste” on which there is no universal, objective truth. Can people increase their chances of selecting options that they will enjoy? One promising approach is to tap into the knowledge of others who have already experienced and evaluated the available options. The recommender systems community has leveraged this source of knowledge to develop *collaborative filtering methods*, which estimate the subjective quality of options for people who have not yet experienced them [81, 1]. One key insight from this field is that, because tastes differ, building recommendations based solely on the evaluations of individuals *similar* to the target individual often improves the quality of the recommendations [53]; similarity is typically defined as the correlation between the ratings of options evaluated by both the target and another person. Although the consumer industry enables people to benefit from recommender systems in some domains (e.g., choosing a movie to watch online), for many everyday decisions neither algorithms nor “big data” are at hand. It thus remains unclear how individuals can best leverage the experience of others when they share prior experience about the available options with only a relatively small community of peers. Should they use strategies that aggregate the opinions of several other individuals or is it better to rely on the opinions of just a few similar others?

Although most everyday decisions are about “matters of taste,” there has been relatively little study of the precise social learning strategies that people can use to inform their decisions. On the one hand, descriptive research in social, consumer, and developmental psychology has shown empirically that people are swayed more by the opinions of similar individuals than by those of less similar others [12, 92, 17, 14, 62, 31, 25]. However, this literature has generally not emphasized precise, formal models of social learning: What exactly are the strategies that people can use? Should everybody use the same strategies and, if not, what are the crucial factors deciding which strategy works best for whom and why? On the other hand, descriptive and normative research on social learning, advice taking, and judgment aggregation in the fields of cognitive science, judgment and decision making, anthropology, and biology is rife with formal models of social learning. However, with a few exceptions [101, 72, 36], these models are about “matters of fact,” where there is an objective ground truth [16,51,48,78,67, 30, 66,9].

In this article, we go beyond the small body of literature on strategies for social learning about “matters of taste” [101, 72, 36] in three main ways. First, we undertake an exercise in *theory integration* by capitalizing on the striking conceptual similarities between seminal recommender system algorithms and both (i) models of judgment and categorization and (ii) models of social learning and social decision making about matters of fact; more specifically, we recast the latter two classes of models as *social learning strategies for matters of taste*. Second, guided by this mapping, we use simulations to investigate how ordinary people with limited experience could use different strategies to leverage the experience of others and thus make better decisions about matters of taste. To this end, we examine the inevitable trade-off between harnessing the apparent (dis)similarity between people’s tastes—to discriminate between more and less relevant people—and estimating those similarities accurately enough on the basis of limited shared experience (i.e., the number of options both the decision maker and the others in the community have experienced). Third, we explore how this trade-off plays out for different individuals depending on how similar their tastes are to those of the other people in the population (e.g., for people with mainstream vs. alternative tastes).

We studied the role of experience and inter-individual differences in taste using a large-scale empirical dataset on people’s ratings of 100 jokes. This dataset has been widely used by the recommender systems community as a benchmark dataset [44]. It is unique in the literature because several thousand people have evaluated all the available options; in other well-known recommender datasets (e.g., MovieLens, LastFM, or Netflix), in contrast, even the most popular items have been rated by only a small fraction of all people in the dataset (i.e., high sparsity). Using such a dense dataset allowed us to treat the ratings of each individual and her peers as a unique prediction environment, in which a given individual’s rating is the ground truth to be predicted. In total, this approach yields 14,000 unique prediction environments with a total of 1,400,00 item evaluations, allowing us to study how the performance of different strategies is affected by the statistical characteristics of each prediction environment. For comparison, the largest testbeds of unique prediction environments for factual problems with informational and social cues contain 63 and 90 environments, respectively [93, 68].

Using this large-scale dataset, we will show that the level of an individual’s simulated previous experience within a domain determines the effectiveness of different social learning strategies. Experienced individuals can benefit from applying strategies that rely on the estimation of similarity. Inexperienced individuals, by contrast, should apply strategies that unconditionally aggregate the opinions of many individuals, despite differences in taste. This result holds across a number of settings (e.g., the size of the community of peers or the average number of items experienced in the population) and for almost all individuals in the dataset. Unless they have a considerable level of experience, people’s estimates of similarity are likely to be error-prone, and relying on them does more harm than good. People with alternative (or less mainstream) tastes and high dispersion in taste similarity with other people could benefit most from accurately identifying similar others—for them, the average experience of the community is typically uninformative. People with mainstream tastes and low dispersion in taste similarity with others, in contrast, are already doing well by unconditionally aggregating the opinions of the crowd. For such people, strategies that rely heavily on similarity cannot beat this simple aggregation strategy—even at high levels of experience.

## 2 Mapping Recommendation System Algorithms to *Informational* and *Social* Cue-Based Strategies

Only a handful of studies have formalized and explored *social learning strategies for matters of taste*, where people use the experiences of others to make predictions about their own future taste experiences, and the conditions under which different strategies thrive or fail. Yaniv et al. [101] studied experimentally the conditions under which people give more weight to the expressed preference of a similar individual than to that of a group of randomly sampled individuals; in a simulation study with synthetic data, they demonstrated the prescriptive appeal of these two simple strategies. Müller-Trede et al. [72] investigated the conditions under which people can benefit from taking the advice of a crowd of similar or randomly chosen individuals, as opposed to focusing on the opinion of just one other individual, sampled either at random or on the basis of similarity. They theoretically studied how people can leverage the advice of several similar others to predict their own future taste experiences and corroborated their predictions using data on people’s preferences for music and short movies. Finally, Gershman et al. [36] proposed a model in which people rely on the revealed choices of other individuals to infer the latent preference groups to which these individuals belong and then use this information to weight the opinions of similar others. In a series of experiments, the authors observed a good correspondence between their model and human behavior.

Here we formalize several social learning strategies for matters of taste and study the extent to which applying these strategies would lead people to choose experiences they will enjoy. We started by identifying conceptual similarities between early recommender system algorithms and models typically applied to the task of predicting matters of fact—that is, where people have access to either *informational cues* potentially related to an objective criterion (e.g., using the number of movie theaters in a city to predict its population size) or *social cues* (i.e., opinions of others on the same objective criterion). We used this correspondence to recast these models as social learning strategies for matters of taste—some of which were inspired by seminal algorithms from recommender system research—that can be used to predict an individual’s future evaluations of an option based on the past evaluations of others.

In the following, we illustrate some of the strategies using the example of deciding which movie to watch based on other people’s ratings. This fictional dataset (see Table 1) has the same structure as the large-scale datasets used in recommender system research and in our own study below. Sofia likes superhero movies and wants to decide whether to watch *Batman* or *Fantastic Four*. Her friends have already seen both movies. She and her friends have all watched and evaluated several other movies. From Sofia’s perspective, her own future taste experiences are the criterion values she seeks to predict, and her friends’ evaluations are cues she can use to predict them. Sofia can leverage the options that she and her friends have evaluated in the past to assess her taste similarity with them (see also [12, 92, 17, 31, 101]), defined as the Pearson correlation between her past evaluations and the evaluations of her friends. Sofia thinks that she and Bob have similar tastes, as they have expressed exactly the same tastes about three options. If Bob truly were her “taste doppelgänger” she could simply imitate his evaluations and arrive at accurate estimates of her own future enjoyment (*doppelgänger* strategy; see Table 2). But it is unclear to what extent this apparent similarity—in just a small set of shared past experiences—will generalize to future cases.

**Table 1:**
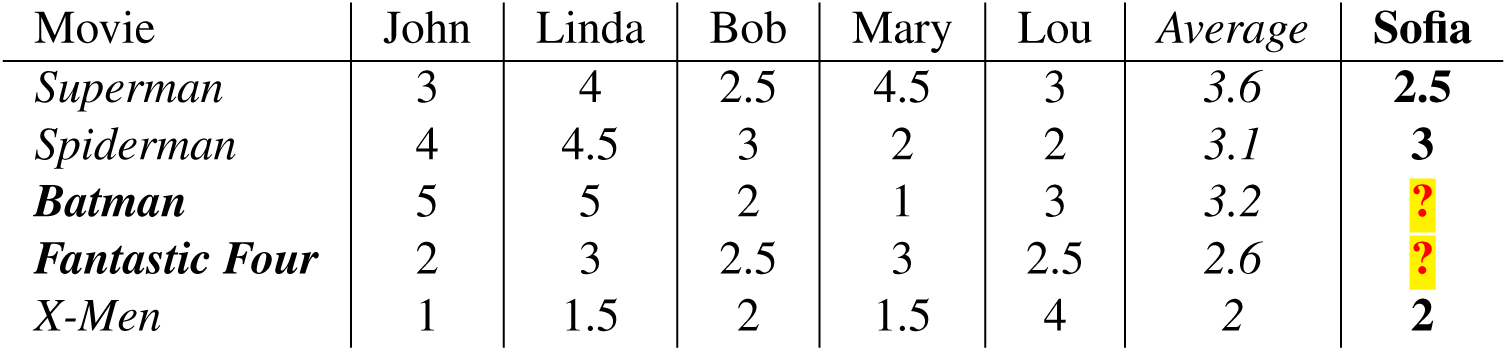
Fictional dataset illustrating the strategies potentially used by humans and recommender algorithms to predict the taste of an individual. *Sofia* is the target individual trying to predict which movie—*Batman* or *Fantastic Four*—she will enjoy more based on the ratings of five friends. The movies are rated on a scale ranging from 1 to 5, where higher values indicate more positive reviews. This prediction challenge is the same both for recommender algorithms relying on collaborative filtering, which predict a user’s taste by considering the ratings of all other users, and for humans who try to predict their own future taste experience based on the experiences of their peers. Some key differences between the two domains are the amount of data available and the respective computational demands: Recommender algorithms operate on very large matrices with thousands of users and items and can do calculations effortlessly, whereas humans are constrained by the limits of their social network and their bounded cognitive capacities.

| Movie | John | Linda | Bob | Mary | Lou | Average | Sofia |
| --- | --- | --- | --- | --- | --- | --- | --- |
| *Superman* | 3 | 4 | 2.5 | 4.5 | 3 | 3.6 | **2.5** |
| *Spiderman* | 4 | 4.5 | 3 | 2 | 2 | 3.1 | **3** |
| ***Batman*** | 5 | 5 | 2 | 1 | 3 | 3.2 | **?** |
| ***Fantastic Four*** | 2 | 3 | 2.5 | 3 | 2.5 | 2.6 | **?** |
| *X-Men* | 1 | 1.5 | 2 | 1.5 | 4 | 2 | **2** |

**Table 2:** Social learning strategies for matters of taste conceptually similar to strategies using informational cues or social cues (i.e., people’s opinions). “−” denotes that we could not find examples for that combination of strategy and research stream in the literature. Strategies incorporating similarity information are typeset in *italics* and those averaging across several individuals’ evaluations are typeset in **bold**. All strategies are person based, except *similar options*, marked in *blue*. All other strategies first estimate the expected utility *û_i_* (i.e., enjoyment) of each option *i* and then select the option with the highest estimated utility; when several options have the same estimated utility, one of the tied options is chosen at random. All the strategies use the Pearson correlation coefficient as a measure of similarity between two individuals *i* and *j*, which is defined as 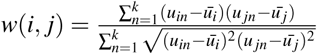. For *similar options*, similarity is calculated using the Pearson correlation coefficient but between two items *k* and *l*.

| Strategy | Social learning strategy |  | Parallels in the literature |  |  |
| --- | --- | --- | --- | --- | --- |
|  | Verbal description | Formal definition | Informational cues | Social cues | Recommender systems |
| *Doppelgänger* [101] | Find individual $s$ with the most similar taste and adopt that individual's evaluations as your own estimates. | $\hat{u}_i = u_s$ | *Take the best* [40], *single attribute* [59] | *Imitate the best, best member* [83, 103] | *Nearest neighbors* ($k = s = 1$) |
| **Whole crowd** [101, 72] | Average the evaluations of all $N$ other individuals (i.e., go with the mainstream). | $\hat{u}_i = 1/N \times \sum_{j=1}^N u_j$ | *Equal/unit weights* [22] | *Averaging* [27] | *Nearest neighbors* ($k = N$), often used as a benchmark [90] |
| *Clique* [72] | Average the evaluations of the $k$ most similar individuals. | $\hat{u}_i = 1/k \times \sum_{j=1}^k u_j$ | – | *Select crowd* [68], *expert crowd* [45] | *Nearest neighbors* $1 < k < N$ [90] |
| *Similar crowd* | Average the evaluations of all $k$ individuals whose taste is correlated with yours above a similarity threshold $t$. | $\hat{u}_i = 1/k \times \sum_{j=1}^k u_j$ | – | – | Common implementation of *nearest neighbors* [23] |
| **Similarity-weighted crowd** | Weight the evaluations of all $N$ individuals according to their similarity to your taste. | $\hat{u}_i = \frac{1}{\sum_{j=1}^N w_j} \sum_{j=1}^N w_j \times u_j$ | *Weighted average* [50, 18] | *Weighted crowd* [20] | *Weighted neighbors* [80] |
| *Similar options* | Find the $k$ most similar options (i.e., with similar evaluation profiles across people) and weight your own evaluations of them according to their similarity. | $\hat{u}_i = \frac{1}{\sum_{l=1}^k w_l} \sum_{l=1}^k w_l \times u_l$ | *Exemplar models* [65, 61], *nearest neighbor models* [3] | – | *Item-based algorithms* [87] |
| Random other [42] | Select an individual $r$ at random and adopt that individual's evaluations as your own estimates. | $\hat{u}_i = u_r$ | *Minimalist* [40] | *Random copying* [15] | Occasionally used as benchmark strategy |

Sofia may thus prefer to take other people’s evaluations into account as well. One approach would be to assign equal weights to all individuals and simply use the average evaluation (i.e., the “mainstream” opinion; *whole crowd* strategy; see Table 2). Yet that would also incorporate the evaluations of individuals with possibly very different—or even antithetical—tastes (e.g., Lou’s ratings seem to be negatively correlated with those of Sofia). To avoid this problem, Sofia could instead rely on the opinions of a select few similar people, such as Bob and Linda (*clique* strategy; see Table 2) or people whose tastes are at least minimally similar to hers (and thus exclude Lou; *similar crowd* strategy; see Table 2). Alternatively, she could assign weights to people’s opinions in proportion to their similarity to hers (*similarity-weighted crowd* strategy; see Table 2). Finally, she could search for movies that the others rated similarly to *Batman* and for movies that they rated similarly to *Fantastic Four*. In line with models of similarity-based inference in cognitive science [73, 47], she could then weight her past evaluations of these proxy movies according to their similarity to the target and choose the one that shows the highest promise (*similar options* strategy; see Table 2; e.g., *Spiderman* could serve as a proxy movie for *Batman*).

The proposed mapping (Table 2) emphasizes the close correspondence between recommendation algorithms on the one hand and informational and social cue-based decision strategies on the other. The social learning strategies can be placed on a continuum between strategies relying solely on similarity information and strategies relying solely on a simple aggregation of opinions irrespective of similarity. Strategies in between those two endpoints put more or less weight on aggregation or similarity, respectively. The mapping reveals groupings and conceptual similarities between the strategies. For example, although the *whole crowd*, *cliqUe*, and *döppelganger* strategies rely on similarity information to a different extent, all three strategies can be seen as instances of the *k* nearest neighbor algorithm from the recommender systems literature [23]; this algorithm relies on a single parameter *k* to determine the number of most similar individuals whose opinions will be aggregated. Crucially, for some of the combinations of strategy and research stream, we could not find any examples in the literature. For example, the *similar crowd* strategy has been implemented in the recommender system literature, but it has not yet been studied as an informational or social cue-based strategy in other domains. Likewise, the *similar options* strategy corresponds to *exemplar models* and *nearest neighbor* models from the informational domain and to *item–item algorithms* in collaborative filtering, but it has not yet been studied in the social-cue domain. These unexplored strategy-domain combinations indicate possibilities for future research and illustrate the value of theory integration in revealing promising new research avenues.

## 3 When and Why Are Social Learning Strategies for Matters of Taste Expected to Perform Well? A Normative Analysis

The performance of strategies using either informational or social cues to predict matters of fact (see Table 2) has been studied extensively in cognitive psychology, forecasting, and machine learning [100, 41, 21, 94, 68]. There are two important insights from these normative investigations. First, the amount of data that is used to train a given strategy is crucial for its performance. Strategies that rely on accurate estimates of many parameters require a decent amount of data before they start paying off [39, 34, 59, 68]. Second, the performance of any strategy depends on the structure of the task environment in which it is applied. Two key factors that typically affect the performance of strategies are the mean correlation between cues and the criterion value (i.e., mean cue-criterion correlation) and the dispersion across those cue-criterion correlations ([68, 93].

Strategies assigning equal weights to all informational cues or averaging the opinions of many individuals tend to perform relatively well when the mean cue-criterion correlation is high and the dispersion in the predictive power of cues is low [27, 59, 68]. Single-cue strategies, which rely only on the seemingly most important cue, tend to perform well relative to more complex strategies when (i) there is a high correlation between the criterion value and the cue deemed most important and (ii) this cue is correlated with the other cues, implying that it already captures the predictive information contained in the additional, less predictive cues [59, 21,20, 68]. Strategies that rely on a few very good predictors or assign different weights to predictors fall in between averaging and single-cue strategies. Those strategies are expected to perform better as the dispersion in the predictive power of the cues in the environment increases because (i) it is easier to identify highly valuable predictors and forecasters [68,71,45] and (ii) differential weighting can profit from differences in the predictive values of the cues [59]. How do these results transfer to the domain of taste?

### Experience—Amount of learning data in a domain

In the domain of taste, the size of the training sample corresponds to the number of options individuals have experienced in the past. Strategies that rely on estimates of taste similarity cannot leverage the knowledge of similar others unless an individual shares enough experiences with those people that she can accurately estimate their similarity. A comparable challenge is faced by recommender system algorithms when recommending options to new users about whom they know little or nothing (the *new User cold-start problem* [29]). Strategies that heavily rely on similarity are expected to improve in performance as an individual acquires new experiences. In contrast, aggregation-heavy strategies, which place little (to no) weight on similarity, should perform well even with little experience because they do not require accurate estimates of similarity. In sum, a large pool of shared experiences between an individual and her peers should lead to better estimation of similarity and give similarity-heavy strategies an edge over aggregation-heavy strategies.

### Similarity to the average person in the crowd—Mean cue–criterion correlation

How does the mean cue-criterion correlation translate to the domain of taste? People’s tastes are subjective. The tastes of each individual in the population can therefore be seen as a unique criterion value that he or she seeks to predict. Because the opinions of others can be treated as predictive cues, it is crucial to know how an individual’s taste correlates with the tastes of her peers. In this setup, the mean correlation between an individual and her peers (i.e., *mean taste similarity*) corresponds to the mean cue-criterion correlation in “matters of fact” problems (see also [72]). Depending on people’s mean taste similarity, the relevance of mis-estimating similarity will vary widely across a population. For people who are, on average, a lot like their peers (i.e., have mainstream tastes), strategies aggregating the opinions of many different peers can be expected to perform quite well, even when experience is limited (see [22, 26, 18]). People with alternative tastes (i.e., mean taste similarity < 0), in contrast, will perform worse than chance by unconditionally following the whole crowd’s opinion. Rather, they can be expected to benefit considerably from finding and relying on just a few similar others.

### Dispersion in taste correlations—Dispersion in the predictive value of cues or experts

For all strategies relying on similarity, the dispersion in the predictive power of the cues is also crucial [63, 68]. Even if the mean predictive power of cues is low, one cue or several cues might still be of appreciably higher quality than the others and capture most of the predictive information in a problem. In the context of matters of taste, such “dispersion in taste similarity” can be operationalized as, for example, the standard deviation of the similarity correlations between an individual and everybody else. Individuals with low dispersion in taste similarity can reap only limited benefits from using similarity-heavy strategies and correctly estimating similarity. For them, any weighting scheme will work equally well (see [22, 18]). People with higher dispersion in taste similarity, in contrast, can learn much more by correctly estimating similarity. For them, assigning the right weights to the right people is crucial. Individuals with low or negative mean taste similarity with others and high dispersion in taste similarity can be expected to benefit the most by correctly assigning weights to other people or by finding a few very similar individuals.

### Regions of best performance

Having established the connection between predicting matters of fact and matters of taste, we can derive predictions about when specific social learning strategies for matters of taste can be expected to perform well: Strategies that rely on the estimation of similarity information should perform better as the level of shared experience increases, whereas strategies ignoring similarity information should have an edge when individuals have little experience in a domain. Yet the level of experience at which people will benefit from switching from aggregation-heavy strategies to similarity-based strategies can be expected to vary considerably depending on their mean taste similarity and the dispersion in taste similarity. For individuals with low mean taste similarity and high dispersion in taste similarity, we expect that little experience is needed before similarity-heavy strategies start outperforming aggregation-heavy strategies. For individuals with high mean taste similarity and low dispersion in taste similarity, in contrast, we expect that more experience is required before similarity-heavy strategies start outperforming aggregation-heavy strategies.

## 4 Simulation Study

To investigate how the amount of experience and individual tastes impact the performance of the proposed social learning strategies (see Table 2), we simulated the performance of those strategies in Jester, a large-scale, real-world dataset [44]. The Jester dataset has been used extensively to study collaborative filtering algorithms. It contains 4.1 million evaluations of 100 jokes by 73,421 participants. In contrast to other datasets studied by the recommender system community, a large number of participants in the Jester dataset evaluated all options. Although other established datasets used in the recommender literature (e.g., MovieLens, LastFM, or Netflix) may contain millions of evaluations, to the best of our knowledge, even the most popular items in those datasets have been evaluated by only a small fraction of all users. Using the rich Jester dataset allowed us to study the principles underlying the success of different social learning strategies at a very large scale [46, 75].

To test our predictions on how the amount of shared experience influences the strategies’ performance, we experimentally varied the number of evaluations used for each of the simulated decision makers (i.e., the number of options previously experienced and rated in that domain; e.g., the number of rows in Table 1; note that the actual participants experienced all 100 jokes, we experimentally manipulated the experience of the simulated decision makers). As simulated experience increases, the strategies relying on similarity could thus base their similarity estimates on more data. Furthermore, the social network from which a person can leverage vicarious experience is likely much smaller than the thousands of people available in typical recommender system datasets. The cognitive limit of the number of stable relationships that people can maintain is estimated to be around 250 [24]. To mirror this real-world feature, we opted to simulate small “communities” of 250 members each (as opposed to letting decision makers have access to all other individuals in the population). In the Supplementary Material, we present variations on the main simulation in which the community size is much smaller (i.e., 25) or equals the entire population (i.e., all 14,000 participants). For all three community sizes, the patterns of results are qualitatively similar and lead to the same conclusions.

### 4.1 Method

The Jester dataset (see http://eigentaste.berkeley.edu) was created by an online recommender system that allowed Internet users to read and rate jokes on a scale ranging from *not funny* (−10) to *funny* (+10) (see also Figure S1 in the Supplementary Material). We used the participants’ funniness ratings both as criterion values and to estimate similarity between individuals.

For simplicity, we only used the data of participants who evaluated all jokes (reducing the number of participants from 73,421 to 14,116). We randomly selected 14,000 participants in order to be able to partition them into evenly sized communities of 250 members each. In line with previous work in the recommender system literature, we used the Pearson correlation coefficient as a measure of similarity between two individuals or two items *i* and *j* [54], defined as 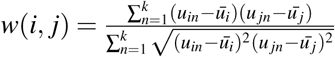. We chose to adopt the Pearson correlation as the measure of similarity so that we could better connect our results to the recommender systems literature, in which it is the canonical measure of similarity.

In each simulation run, we performed the following steps:

- From the 14,000 individuals, we randomly generated 56 communities with 250 members each (14,000/250).
- We randomly divided the jokes into a training set (*x* jokes) and a test set (25 jokes); this assignment was the same for all individuals within a particular community but differed across communities. For each individual within each community, the different social learning strategies were fitted to the training set, assuming that individuals could only access the ratings of their peers within their own community.
- For each individual (within each community), we generated all 300 possible pair comparisons within the test set (*K* × (*K* − 1) ÷ 2, where K stands for the number of items in the test set). For each strategy, we recorded which of the two jokes in a pair the strategy predicted would be rated higher by that individual.In the very few cases where two jokes had exactly the same predicted or actual funniness rating, ties were broken at random.
- For each strategy and level of experience (*x* = 5, 10 … 70, 75), we recorded the performance of each strategy, that is, its proportion of correct predictions across the 300 pair comparisons for each individual. Formally, performance can be expressed as *C* / (*K* × (*K* − 1) ÷ 2), where C stands for correct predictions.

This procedure was repeated 2,000 times. It yielded 600,000 pair comparisons per individual (300 × 2,000), 150,000,000 pair comparisons per community (300 × 250 × 2,000), and 390,000,000,000 pair comparisons in total (30,000 × 250 × 56 × 2,000). To derive the final results, we averaged across the 2,000 repetitions for each combination of individual, level of experience, and strategy. We identified the best performing strategy for each individual based on the average performance of each strategy (i.e., averaged across the simulation runs).

We investigated how the strategies’ performance changed as a function of simulated experience by repeating the procedure for different numbers (*x*) of jokes experienced in the training set (from 5 to 75 in steps of 5).In the Supplementary Material, we also present the results for several variations of this baseline simulation, all of which yielded similar qualitative results and the same conclusions. The standard errors of the performance measure ranged between 0.00071 and 0.00287 across all individuals, experience levels and strategies.

### 4.2 Results

We start by looking at the average performance across all 14,000 participants and examining how the different strategies perform as a function of simulated experience (i.e., the number of jokes used to estimate similarity). Going one step further, we then investigate how the strategies perform for different individuals, depending on how their tastes correlate with those of others. We examine whether an individual’s mean taste similarity with other participants and dispersion in taste similarity impact the success of different social learning strategies as predicted. The performance of a strategy is defined as the proportion of correct predictions in all the pair comparisons generated in the test set and averaged over repetitions of the simulation (see subsection 4.1). Figure 1A shows the average performance of each strategy (across individuals) at different levels of experience; Figure 1B shows the proportion of individuals for whom a given strategy performed best at those different levels of experience.

**Figure 1:**
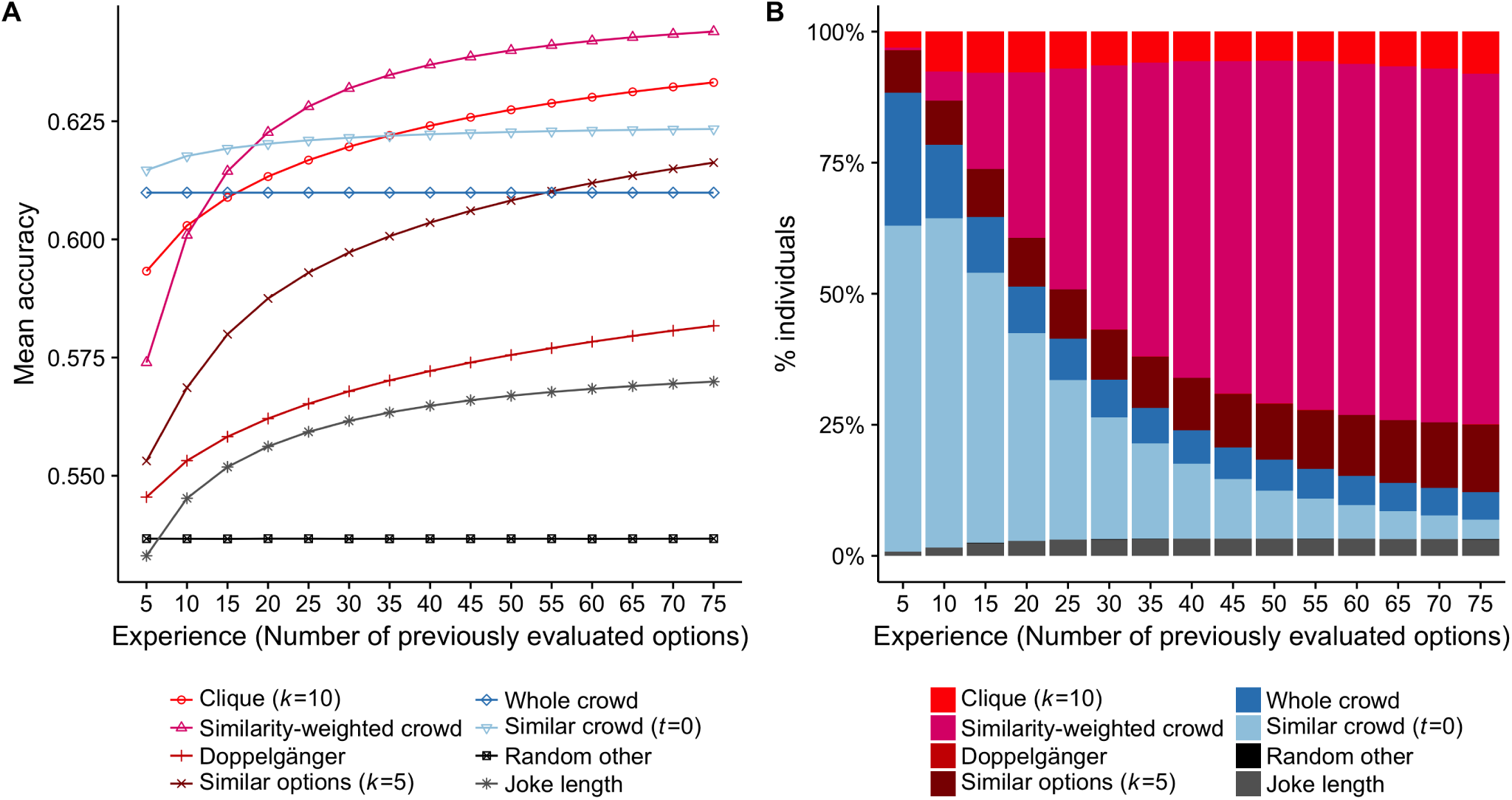
**Panel A:** The performance of social learning strategies as a simulated decision maker’s experience increases (i.e., number of jokes experienced and evaluated). The curves show the average performance across all 14,000 participants. The strategies are grouped by color into those that rely primarily on aggregation (blue), those that rely primarily on similarity (red), and three benchmark strategies (i.e., the *random other* strategy, in black, see also Table 2; a *joke length* strategy, which uses the length of the joke to infer its quality, in grey; and the *similar options* strategy, which corresponds to an item-item collaborative filtering algorithm and uses people’s ratings for the *k* most similar options as a proxy to decide which of the two unevaluated options to choose, in brown; see also Table 2). The *random other* strategy predicted approximately 54% of pairs correctly, which indicates that there is a small shared sense of humor in the population (i.e., slightly better than chance; see also Figure S8 in the Supplementary Material). **Panel B:** The percentage of simulated decision makers for whom each strategy performed best (y-axis) as a function of the number of options experienced (x-axis). The *doppelgänger* and *random other* strategies are barely visible because they almost never performed best for any participant at any level of experience.

#### 4.2.1 How does the strategies’ performance change as experience increases?

Although strategies that use a combination of similarity and aggregation perform best when decision makers have experienced many options (Figure 1A), this is not the case at low levels of experience. For the lowest levels of experience, following either the *whole crowd* or a *similar crowd* is the best performing strategy. Following a *similarity-weighted crowd* or a *clique* starts outperforming the *whole crowd*, which aggregates opinions unconditionally, only after approximately 15 options have been experienced. The *similarity-weighted crowd* strategy outperforms the *similar crowd* strategy after approximately 20 options; the *clique* strategy needs to experience 35 options to do so. These aggregate results are corroborated by individual-level analyses (i.e., proportion of decision makers for whom a strategy performed best; Figure 1B). At the lowest level of experience, the aggregation strategies perform best for more than four fifths of the population. As decision makers become more experienced, the proportion of the population for which these strategies perform best wanes. At the highest level of experience, they perform best for only 10% of the population. Overall, the performance of strategies that heavily rely on similarity improves markedly as experience increases, whereas strategies that rely more on aggregation (and less on similarity) start out at high performance levels—even with little (or no) experience—but improve only little, if at all.

#### 4.2.2 The informational value of dissimilar others

To sum up the results so far, decision makers who have not yet experienced many options are well advised to simply aggregate the evaluations of individuals who seem to have at least minimally similar (i.e., positively correlated) tastes (*similar crowd*) or even to unconditionally aggregate the evaluations of all individuals (*whole crowd)*. Although the opinions of truly similar individuals are more informative than those of truly dissimilar individuals, relying more on seemingly similar individuals is only beneficial to the extent that the similarity estimates are accurate enough. When experience is limited, estimates of similarity are apparently often not accurate enough to be of much—or any—use. The *doppelgänger* strategy (i.e., relying solely on the most similar people) does not perform that well because of the difficulty of reliably estimating similarity based on limited experience. Mirroring results from research on the wisdom of *small* crowds [45, 68], our findings showed that taking into account additional—although less similar—peers and averaging their recommendations markedly improves performance. This point can be seen in the *clique* strategy, where the number of peers (*k*) whose evaluations are averaged determines how selectively or broadly this strategy relies on similarity. In the results presented in Figure 1, *k* was fixed at 10. Additional simulations that varied the value of *k* showed that with little experience it is better to rely on large cliques (approximately 100), whereas with the highest level of experience, performance peaks with moderately sized cliques (approximately 30; see Figure S8 in the Supplementary Material).

#### 4.2.3 Interindividual variability in tastes

So far we have examined how strategies perform irrespective of the interindividual differences in how people’s tastes correlate with those of other people. Yet, as outlined in the previous section, the cost of mis-estimating similarity can be expected to be largest for people whose tastes differ markedly from their peers’ average tastes and whose peers differ considerably among themselves in how similar they are to the target person.

To investigate the relation between the statistical structure of taste and the strategies’ performance, we first need to quantify these two aspects of people’s tastes. To this end, we calculated for each of the 14,000 individuals the mean and standard deviation of their taste correlations with all 13,999 potential peers, indicating the average similarity with their peers and the dispersion in those taste similarities, respectively. The overall average similarity and dispersion were *μ* = 0.11 and σ = 0.13, respectively, which indicates a relatively low level of shared taste in the population. By statistical necessity, the tastes of the majority of people are positively correlated with those of other individuals, yet a sizable minority of individuals have neutral—or even antithetical—tastes when compared to the mean evaluations of the entire population (as reflected by near zero or negative mean correlations; see Figure 2). On both sides of this spectrum, we can observe individuals with both high and low dispersion of taste similarity across peers (as indicated by the standard deviations). To put the observed values in context, we can contrast them with those of two idealized, synthetic individuals. First, a perfectly mainstream individual whose tastes are identical to the predictions of the *whole crowd* (i.e., whose evaluation of each joke is identical to the average evaluation across all participants) would have a mean taste similarity of 0.33 and a dispersion of 0.18. Second, an idiosyncratic individual whose appreciation of the jokes is random (i.e., sampled from uniform distributions covering the whole range of the evaluation scale) would have a mean taste similarity close to 0 and a dispersion close to 0.08. This comparison shows that people differ markedly in how similar their tastes are to those of their peers and that every individual thus represents a unique environment for contrasting the performance of the social learning strategies.

**Figure 2:**
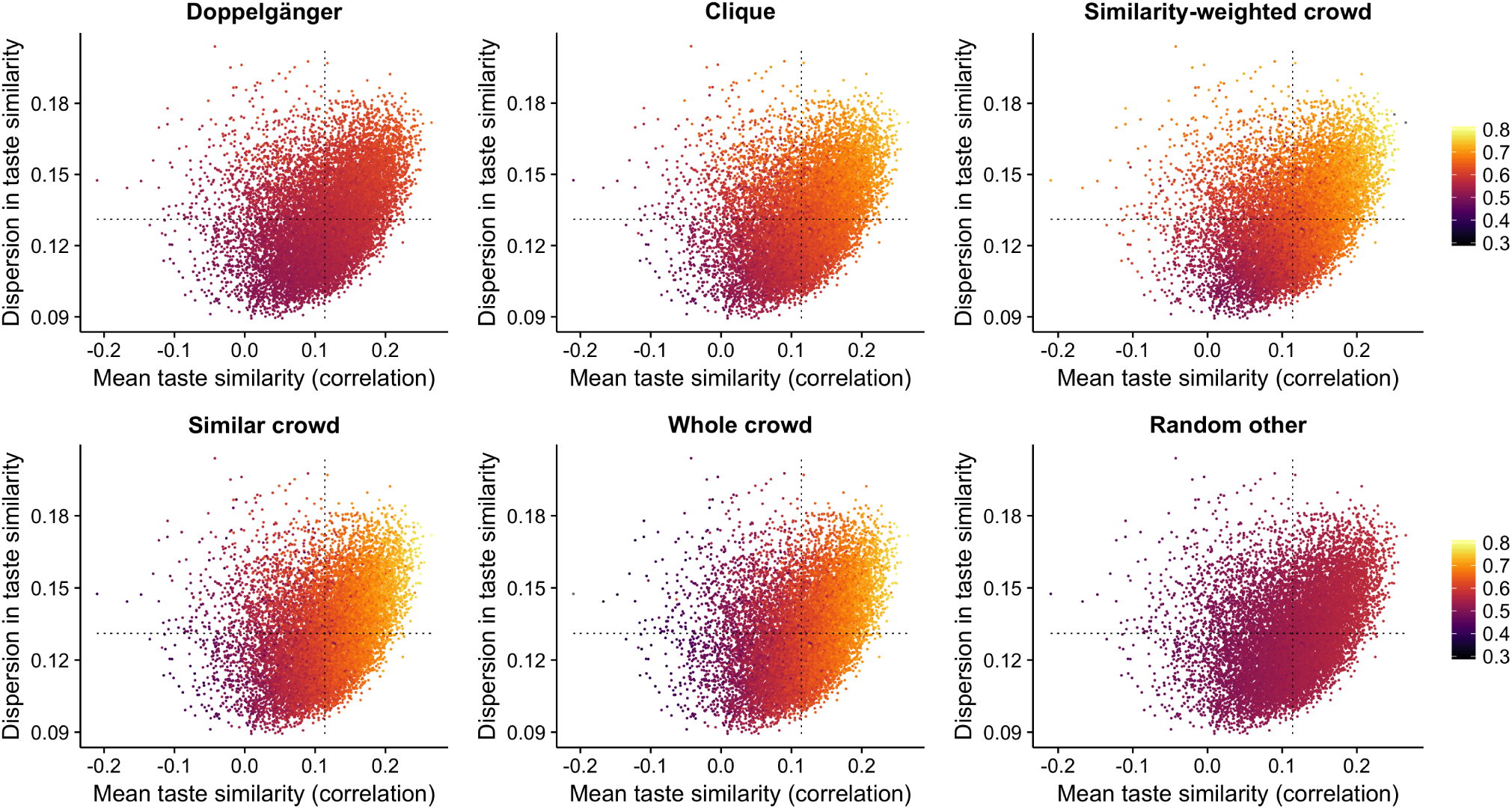
Strategies’ performance depending on how people’s tastes correlate with those of others in the population (after experiencing 25 options). The scatter plots represent the strategies’ performance for each of the 14,000 individuals (i.e., proportion of correct predictions, color coded). Each point represents one of the 14,000 individuals and thus represents a unique prediction environment for social learning strategies for matters of taste. Each individual is positioned according to their mean taste similarity with all other 13,999 individuals (x-axis) and the dispersion in taste similarity with other individuals (i.e., standard deviation of these correlations; y-axis). Dashed horizontal and vertical lines show the overall average of the mean correlations (0.11) and standard deviations (0.13). All strategies perform better for individuals with more mainstream tastes and higher dispersion in taste similarities with their peers except for *random other*, which randomly selects a person to imitate by design. Results are based on averaging across 2,000 repetitions of the simulation. The standard errors of the performance measure across all individuals, experience levels and strategies ranged between 0.00071 and 0.00287 with a median of 0.0028.

#### 4.2.4 How well do the strategies perform for different individuals?

Figure 2 shows how well the strategies perform for individuals with different tastes, ranging from mainstream to alternative, and with different dispersions in taste similarity. For illustration, we focus on the level of experience at which the best-performing similarity-heavy strategies and the aggregation-heavy strategies perform similarly well (approximately 25 options; see Figure 1) and highlight the interaction between mean taste similarity and dispersion in taste similarity (for results at other levels of experience, see Figure S10 in the Supplementary Material). First, as expected, all strategies perform best for individuals with high mean taste similarity. The *whole crowd* strategy correctly predicts more than 70% of the choices for individuals with a high mean taste similarity, but performs worse than chance for individuals with alternative tastes. Crucially, for all strategies that rely on similarity to some extent, this result is moderated by dispersion in taste similarity, as comparing two individuals with the same mean similarity shows (see Figure 2): As we move towards lower mean similarity, the losses in predictive ability are smaller for individuals with high (as compared to low) dispersion in taste similarity with their peers. This result is modest for the *similar crowd* strategy, which only excludes dissimilar individuals, and for the *doppelgänger* strategy, which relies on only the most similar individual, but it is particularly pronounced for the *similarity-weighted crowd* and *clique* strategies, where the differences in performance are as big as 10 percentage points (to verify this, follow a vertical line from bottom to top in Figure 2).

#### 4.2.5 Which strategies performed best for different individuals and levels of experience?

Figure 3 shows the best performing strategy for each individual for three levels of experience: people who are relatively inexperienced (10 experiences), moderately experienced (25 experiences), or who have experienced all options in the training set (75 experiences). Among inexperienced decision makers, the aggregation strategies perform best for almost all individuals with a positive mean taste similarity. A comparison of the aggregation-heavy strategies shows that the *whole crowd* strategy dominates for individuals with high mean taste similarity and low dispersion in taste similarity, while the *similar crowd* strategy performs best for individuals with moderately high mean taste similarity and high dispersion in taste similarity. The strategies that rely on similarity perform best only for individuals with alternative or neutral taste similarity and high dispersion in taste similarity. The pattern of results for moderately experienced decision makers is very different. Here, aggregation-heavy strategies outperform similarity-heavy strategies only for individuals with positive mean taste similarity and low dispersion in taste similarity. The *similarity weighted crowd* strategy performs best for individuals with high dispersion in taste similarity, while the *clique* strategy performs best for individuals with low or negative mean taste similarity and low dispersion. At high levels of experience, the similarity-heavy strategies take over most of the occupied parameter space. At such high levels of experience, the *whole crowd* and *similar crowd* strategies still perform best only for a pocket of individuals with high positive mean taste similarity and very low dispersion in taste similarity.

**Figure 3:**
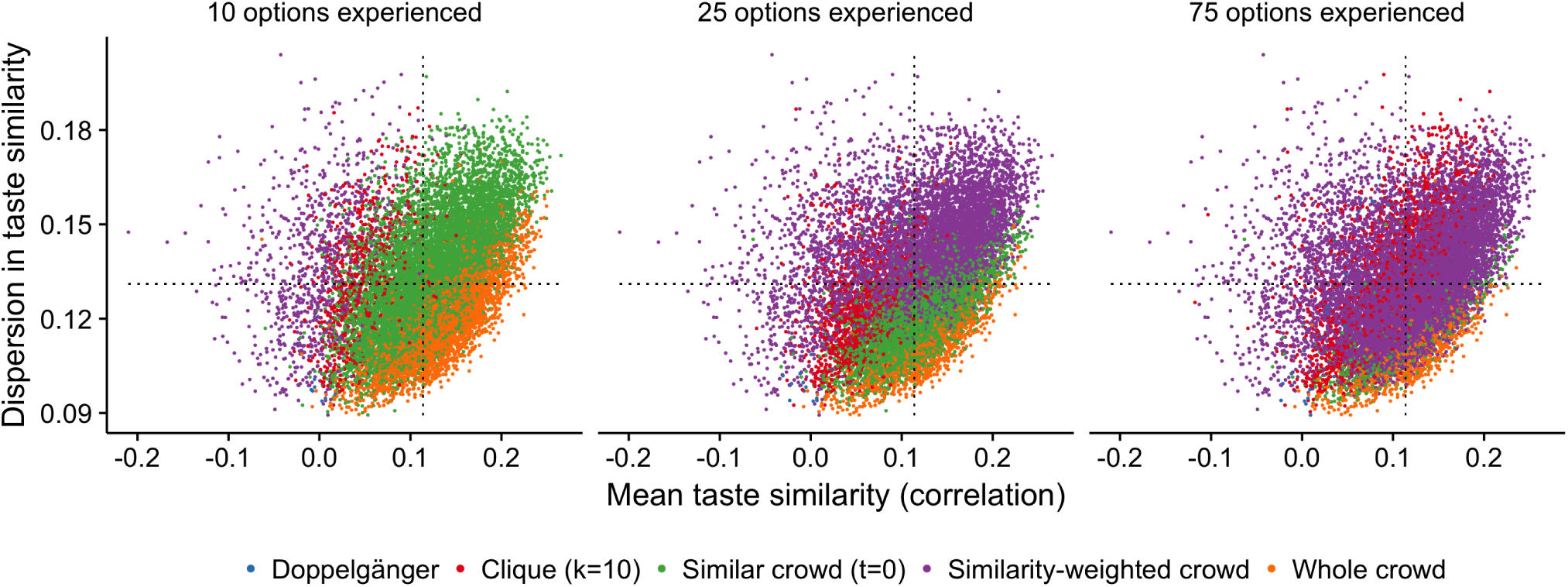
Best-performing strategy depending on how people’s tastes correlate with those of others in the population. The scatter plots show for each individual the best performing strategy when only 10, 25, or 75 random items, respectively, were used to estimate similarity (left- to right-most panel; 75 items represents the highest level of experience used in the simulation). Each point represents one of the 14,000 individuals and thus a unique prediction environment for social learning strategies for matters of taste. Each individual is positioned according to its mean taste-similarity with all other 13,999 individuals (x-axis) and the dispersion in taste similarity with other individuals (i.e., standard deviation of these correlations; y-axis). Dashed horizontal and vertical lines show the overall average of the mean correlations (0.11) and standard deviations (0.13). Results are based on averaging across 2,000 repetitions of the simulation. Standard errors for all individuals, strategies, and experience levels ranged between 0.00071 and 0.00287 with a median of 0.0028.

## 5 General Discussion

Whether it is a matter of which massive open online course (MOOC) to enrol in, which music album to download, or which clothes to buy for the next summer season, most decisions made in everyday life are about matters of taste. There is no unanimous verdict on whether Cornell offers a better statistics course than NYU, on whether John Coltrane’s *Giant Steps* album is better than David Bowie’s *Station to Station*, never mind on styles of clothing [60]. In this article, we set out to understand why some social learning strategies might work for some people but not for others. We show that individuals’ past experiences and the way their tastes relate to those of others interact and jointly determine the effectiveness of different social learning strategies.

### 5.1 Reconnecting Cognitive Science and Recommender Systems

From Thurstone’s discrimination theory [96] to Rosenblatt’s perceptron [85], psychological theories have often influenced the development of new statistical and predictive tools. Inversely, new statistical tools, such as regression analysis and signal detection theory, have inspired the development of new psychological theories [38, 4]. Somewhat surprisingly, early work on the wisdom of crowds and opinion aggregation in psychology [49, 58, 97] seems to have gone unnoticed by recommender systems researchers, although the first algorithms they developed were very similar to these strategies in both spirit and content. Likewise, insights emerging from recommender system research in the last two decades have not really been incorporated back into psychology and the behavioral sciences more generally, although the recommender system community has maintained a general interest in using insights from the behavioral sciences [8, 28] (and many recommender systems researchers suggest that the origins of recommender systems research can be traced back to cognitive science [82]). It is a historical vagary that cognitive psychologists and behavioral scientists have not forcefully addressed the issue of individual learning from the experience of similar others given that they have developed very similar models in the domain of facts.

A key reason for the divergence between these two strands of research is the lack of large-scale datasets amenable to psychological study and interpretation. The recommender datasets leveraged in industrial applications (e.g., Netflix, LastFm, or Pandora) or developed under the auspices of research institutions (e.g., MovieLens) are typically very sparse. Even the most prolific users have evaluated only a small subset of all available options and even the most popular options have been rated by only a small fraction of all users. As a result, researchers have to deal with the substantive challenges introduced by missing data (e.g., by adding values artificially to the matrix [86] or introducing algorithms that cope with missing data [88]), and comparisons between individuals become cumbersome. In this paper, we used the only full large-scale recommender dataset known to us (i.e., with a substantial number of items evaluated by everybody in the population). Additional full datasets from other domains of experience could facilitate cross-fertilization between behavioral scientists and the recommender systems community. Future dialogue between these disciplines is crucial for understanding the social learning strategies that individuals can use to harvest other people’s experiences and thus to inform their own choices in matters of taste.

### 5.2 Experience and the Bias–Variance Trade-Off

With increasing experience in the domain, the predictive ability of all the best-performing strategies increased— except for the strategy relying on the wisdom of the *whole crowd*, which unconditionally averages across all people and is—by design—unaffected by the increasing accuracy of the similarity estimates. All strategies lie on a bias–variance continuum (see Figures S12 and S13; in the Supplementary Material, we present detailed results and an extended discussion of the bias–variance trade-off that corroborate the claims we make below). At one extreme, the *whole crowd* strategy assumes that everybody has the same taste and performs well for individuals whose tastes are indeed well aligned with those of other people. From a bias–variance trade-off perspective [35, 39, 11, 37], this strategy suffers from potentially high bias, especially for people with alternative tastes, but it exhibits zero variance in its prediction error because it does not estimate any free parameters and makes the same prediction regardless of the past experiences an individual has acquired. Strategies relying on similarity, in contrast, have a comparatively low bias because they can adapt to the homogeneity or heterogeneity of tastes in the population. However, they potentially suffer from variance because their predictions depend on the training sample used to estimate similarity. The predictions differ because they are tuned to the sample of experiences that was used to train the model, but relying too much on a particular sample can lead to over-fitting. At the other extreme of the bias–variance continuum is the strategy of adopting the evaluations of only the seemingly most similar person. This strategy potentially allows people to profit from the experiences of their taste *doppelgänger* but it is most reliant on an accurate estimation of similarity and thus most vulnerable to variance.

Each individual in the dataset has her own bias–variance profile (see Figure S14), which depends on how her taste is correlated with that of the other people (see Figure 2 or 3). For most of the strategies (e.g., *similar crowd* and *doppelgänger*), the bias component of the error term differs much more across individuals than the variance component. Apart from the *whole crowd* strategy, which is a zero variance strategy for everybody by definition, there are only mild interindividual differences in the variance component of the error term. The *similarity-weighted* strategy is an exception to this general observation. Although it suffers, on average, from only moderate variance, its variance is much larger for individuals with low or even negative mean taste similarity with other people than for individuals with more positive mean taste similarity. In the former case, the success of the strategy depends strongly on the accuracy of the weights assigned to the few truly very similar individuals. In contrast, if all peers are very similar to the target individual, it matters less what weight is assigned to them.

In sum, these bias–variance results indicate that social learning strategies as well as recommendation algorithms should be evaluated at the individual level, rather than the aggregate level, which has thus far been the common practice in the recommender systems literature. Furthermore, our bias–variance results emphasize the importance of accounting for the differing flexibility of social learning strategies with respect to how well they can tune themselves to past experiences, and in how this flexibility is then revealed in the bias–variance trade-off. To this end, modelers can rely on the established toolbox of model selection techniques (such as cross-validation [5], structural risk minimization [98], or information criteria, such as AIC, BIC, or MDL [2, 89, 84, 77]).

### 5.3 From Matters of Fact to Matters of Taste

Each individual in our study represents a unique prediction problem. Some people’s tastes are very similar to those of their peers; others have opposing preferences. At both ends of this spectrum, people’s taste similarity with other people can be more or less dispersed. To what extent were our predictions about the absolute and relative performance of various social learning strategies for different individuals borne out, and how do our findings differ from those of studies investigating strategies for “matters of fact”?

A number of similarities and differences stand out. First, we found that when experience is scant, strategies relying heavily on averaging performed best for all individuals with a positive mean taste correlation with the crowd. This finding is in line with results on matters of fact showing that, for small sample sizes, equal weighting of cues outperforms differential weighting models [59, 94]. The *similarity-weighted crowd*, which was the best performing similarity-heavy strategy, started to outperform averaging strategies once people had acquired some experience. Yet the exact amount of experience depended both on the mean taste similarity between an individual and the others and on the dispersion in those taste similarities–and for people with high mean taste similarity and low dispersion this never happened (see Figures S4A and S4B in the Supplementary Material). The number of experiences required for similarity-heavy strategies to outperform aggregation strategies might be even larger in real-life settings where people commonly associate themselves with individuals similar in terms of demographics, values, and cultural tastes—a tendency widely documented across societies and contexts and referred to as homophily [70, 32]. Then, people might not need to discount the experiences of dissimilar others as such dissimilar people will not be part of their communities anyway. The prediction problem people face in many real life problems might thus be closer to those faced by the individuals with high mean taste similarity and low dispersion in our simulation (see the lower right quadrant of Figures 2 and 3). With small differences in taste, a wealth of experience is needed to reliably exploit those small differences and to then outperform the simple aggregation strategies.

The *clique* strategy, which averages the evaluations of the 10 most similar individuals, was consistently among the best-performing strategies for experienced individuals; for people with alternative tastes, it was the best-performing strategy even with only scant experience (this result was also replicated in much smaller communities of individuals; see Figure S7B in the Supplementary Material). For individuals with mainstream taste, however, several dozen experiences were required for the *clique* strategy to outperform aggregation-heavy strategies like the *similar crowd* strategy (see Figures S3D and S4A in the Supplementary Material). Overall, our results corroborate findings on the potential of select crowds to solve prediction problems [68,45]. However, they also show that in the domain of taste, much more experience might be required for the potential of small crowds to be realized.

Studies investigating environments with only a small number of cues, either informational or social, have shown that the strategy of relying on just the most predictive cue can perform on par with strategies combining multiple cues [94, 59, 101]. In our study, for experienced individuals with antithetical tastes, the *doppelgänger* strategy (i.e., imitating the seemingly most similar person) performed better than the *whole crowd* strategy, but the *doppelgänger* strategy was almost always dominated by the *clique* and the *similarity weighted crowd* strategies (for some comparisons with other strategies, see Figure S3 in the Supplementary Material; see also [68]). It has been argued that the commonly observed superior performance of cue-based heuristic strategies, which rely on a few pieces of information or ignore cue weights, can be attributed to their lower variance relative to more complex strategies [39]. Indeed, this was true for the people for whom the *whole crowd* strategy performed best (i.e., a strategy ignoring cue weights). In contrast, it did not hold for the few people for whom the single-cue strategy for predicting matters of taste (*doppelgänger)* performed best. This strategy had the highest variance for almost everybody in the population (see Figure S12 in the Supplementary Material). This discrepancy with the corresponding results from research on matters of fact can be attributed to two factors. First, because every individual is a potential cue for every other individual in matters of taste, the number of cues (i.e., number of peers in a community—250 in our simulation) is larger than the number of observations and is much larger than the number of informational or social cues typically studied in matters of fact, where there typically are many more observations than cues. It is well-known that such a low ratio of observations to predictors results in high variance for unbiased strategies [52]. Second, in our study, many cues have approximately the same predictive value, which limits the usefulness of differential weighting or even of selecting the single-best cue or person (as in the *doppelgänger* strategy). In studies on matters of fact, in contrast, the predictive power of the informational cues has typically differed greatly [93].

### 5.4 Model-Based, Content-Based, and Hybrid Social Learning Strategies

In most domains of everyday experience, people (and machines) have access to information beyond their own and other people’s past experiences: informational cues describing options (i.e., features such as a movie’s genre) and other individuals (e.g., a person’s clothing style or personality). The use of such information has been examined extensively in multiple-cue judgment and categorization learning in cognitive science, in content-based and demographic-/personality-based recommender systems [7, 1, 76, 74, 33] and, more generally, in supervised learning in machine learning. People may use these cues (i) to take advantage of the predictive information in the options’ features (e.g., it is a superhero movie) or (ii) to improve their assessment of similarities with their peers (e.g., by considering their age, gender, personality, profession, or social network) [14, 43, 101, 99, 25]. Information going beyond shared experiences might be particularly beneficial when people make their first choices in a domain that is new to their network— for example, when looking for a restaurant in a city they have never visited before. In such situations, when they lack shared experiences with similar others, using the option’s cues to predict its quality directly or using cues characterizing another person (e.g., clothing) to assess similarity could prove beneficial.

But how do people learn from others when they have access to more than one type of information or model? Both the recommender systems literature and the psychological literature on inferences about “matters of fact” are rife with ideas on how to combine different sources of information or models [10, 13, 55, 57, 56]. In the recommender systems context, a well-known example of aggregating different sources of information and approaches is the Netflix competition, in which the winning model was a hybrid of several best-performing models [6]. People, by analogy, could regard learning from similar others and learning from the features of the objects as two modules that operate independently and can be aggregated or used in a complementary manner when a judgment needs to be made. The study of the theoretical performance of hybrid decision strategies, and their prescriptive and descriptive value as models of social learning for matters of taste, are fruitful avenues for future research. Müller-Trede et al. [72] have made inroads in that direction by examining whether people could benefit from combining their own predictions about whether they will like an option (on the basis of prior information about the available options) with the predictions of a crowd of other people. In the language of recommender systems [13], their approach would correspond to a hybrid model between a content-based system, which can derive predictions based on the features of the considered options, and a collaborative system, which relies on the wisdom of similar others.

### 5.5 Learning About Oneself vs. Learning About Others

People can use the strategies investigated in this paper not only to learn about themselves, but also to learn about other people’s preferences. Our theory integration revealed that—from an algorithmic perspective—learning about others, as recommender systems do, and learning about oneself are equivalent problems. Instead of comparing their own taste with that of their friends, individuals can use other people’s past evaluations or other sources of information to assess similarity in tastes between a target individual and other people (e.g., my friend Bob has similar music tastes with John but less so with Linda). Learning what other people like is an invaluable social skill. Thus, it comes as no surprise that toddlers already have an awareness of other people’s preferences [79]. As adults, humans learn about their friend’s preferences; as parents, they learn about their children’s likes and dislikes [69]; and it is good for people to know what their romantic partners like [19]. Being able to better predict other individuals’ tastes can lead to better advice (e.g., recommending a restaurant) and better surrogate choices when picking an option on their behalf (e.g., choosing a dish at a restaurant) [91]. Tapping into people’s knowledge of the preferences of their friends or partners, a few studies have compared the prediction performance of human recommenders with that of actual recommender systems [95, 64, 102]. The results have been inconclusive so far, suggesting that in some contexts humans provide better recommendations than algorithms, but in other contexts recommender systems outperform humans. In addition to prediction comparisons, future research could characterize human and artificial recommenders in terms of comparable formal learning strategies. This may help to clarify the informational or computational advantages that give an edge to humans or machines and open up new avenues for research.

## 6 Conclusion

From picking a restaurant to choosing a movie, most choices people make are on “matters of taste,” where there is no universal, objective truth about the quality of the options available. Still, people can learn vicariously from the experiences of individuals with similar tastes who have already experienced and evaluated options—as exemplified by recommender systems. Mapping out the conceptual similarities between seminal recommender system algorithms, on the one hand, and models of judgment and decision making (based on informational or social cues), on the other hand, we recast the latter as social learning strategies for matters of taste. Furthermore, drawing on research from judgment and decision making, forecasting, and machine learning, we predicted which strategies would perform better or worse for different people, depending on the structure of individuals’ tastes and how much experience they have in the domain. We ran computer simulations on a large-scale, empirical dataset, and investigated how people could leverage the experiences of others to make better decisions. We showed that experienced individuals can benefit from relying mostly on the tastes of seemingly similar people. Inexperienced individuals, in contrast, cannot reliably estimate similarity and are better off picking the mainstream option despite differences in taste. We found that the level of experience beyond which people should switch to similarity-heavy strategies varies substantially across people and depends on (i) how mainstream (or alternative) an individual’s tastes are and (ii) the level of dispersion in taste similarity with the other people in the group. It is a truism that you cannot argue about taste. Nevertheless, we have shown that *which* social learning strategy for matters of taste works best and *for whom* is not subjective—rather, it is subject to rational argumentation.

## 7 Data and code availability

Data and code can be accessed at: https://osf.io/tj7xz/?view_only=74e7c4f81fc046deaee5b1ffafbe8860

## Acknowledgments

We thank Charley M. Wu, Johannes Müller-Trede, David Wolpert, Tobias Schnabel, Dan Cosley, and Thorsten Joachims for their insightful comments. We are indebted to the members of the ABC and ARC research groups at the Max Planck Institute for Human Development, Berlin, and the participants in the Microsoft Artificial Intelligence seminar at Cornell University for their constructive feedback during presentations. We are grateful to Anita Todd and Susannah Goss for editing the manuscript. This research was supported in part through NSF Award IIS-1513692.

## 8 Author Contributions

P.P.A, D.B, and S.M.H. conceived the research, developed the simulation framework for the main results and the bias–variance decomposition, analyzed the simulation results and wrote the paper.

## 10 Supplementary Material

### 10.1 The Jester dataset

#### 10.1.1 Rating Interface

**Figure S1:**
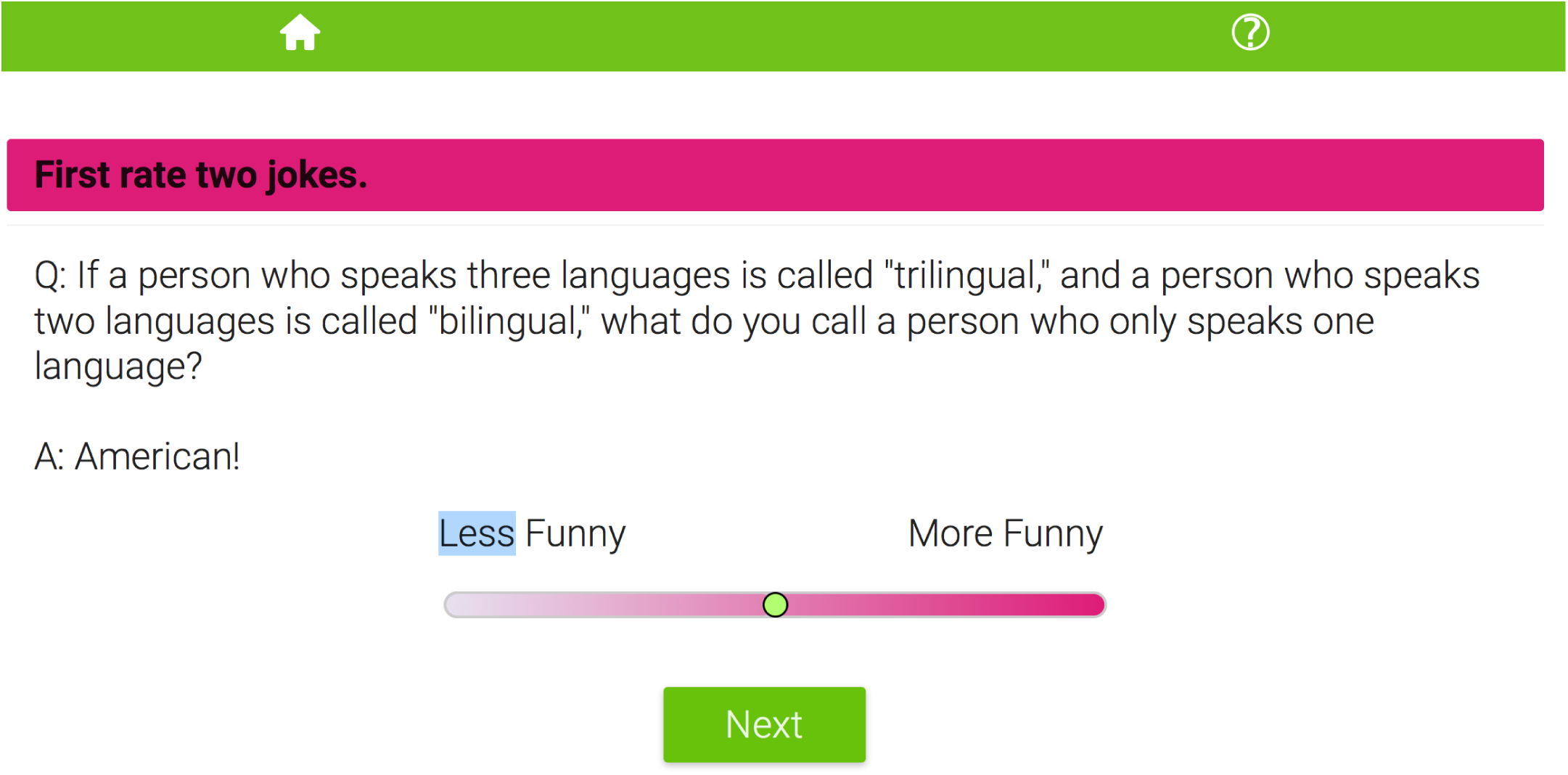
A screenshot from the Jester interface (retrieved on 29 April, 2017).

The first version of the Jester interface was deployed in 1999 by Ken Goldberg, Dhruv Gupta, Hiro Narita, and Mark DiGiovanni at Berkeley University. To the best of our knowledge, the Jester dataset is the only recommender dataset where a large number of participants (14,116 individuals) have evaluated all of the curated options. The interface design is straightforward and has not changed over the years other than for color brush-ups. The jokes appear at the top of the screen. Users can move a continuous slider located below each joke to rate it, from “less funny” at the leftmost end of a scrollbar to “more funny” at the rightmost end. The slider is initially placed in the middle of the scrollbar, corresponding to a neutral joke evaluation score of 0. Moving the slider to the left yields negative ratings; moving it to the right yields positive ratings. Possible ratings range between −10 and 10. At any point, the user can press the “next” button, which is located beneath the scrollbar. Users can read and rate up to 100 jokes. After a random sample of jokes has been evaluated, the system starts recommending jokes to users until the pool of 100 jokes is exhausted.

#### 10.1.2 Average Quality of the Jokes as Reflected by Their Ratings

How much did the jokes differ in terms of their average quality, as indicated by the participants’ ratings? Because strategies such as the *whole crowd* rely exclusively on population mean ratings, it is important to see how much variability there is in the ratings. Figure S2 shows that the “best joke” received an average rating of 3.82, while the “worst” scored −3.43. Ninety-eight of the jokes had unimodal distributions. Two jokes polarized the audience, as reflected by bimodal distributions that peaked on both edges of the evaluation scale (jokes 71 and 75). For many jokes the mode of the ratings is 0, which suggests an anchoring effect of the initial placement of the slider at the middle of the bar.

**Figure S2:**
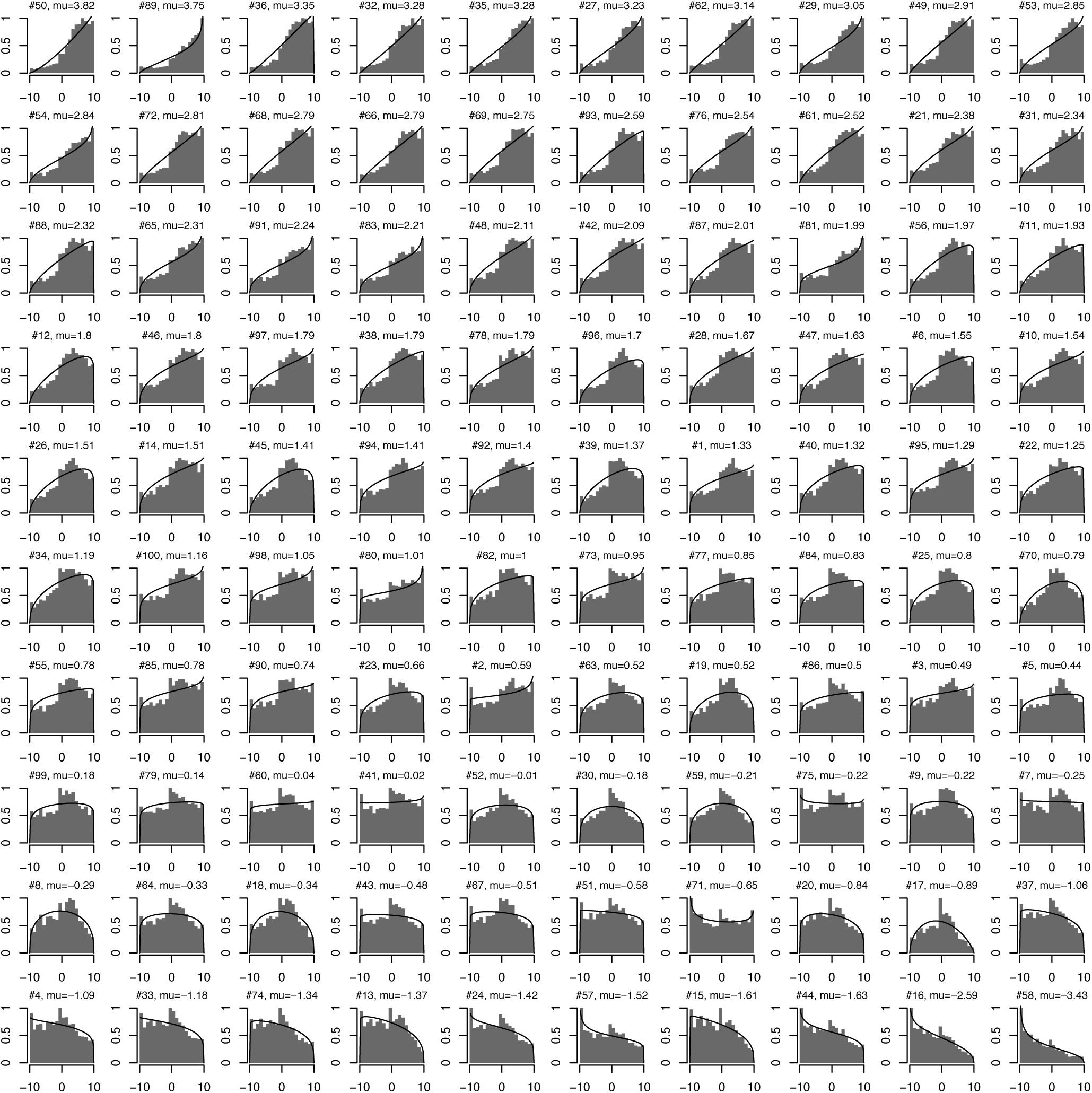
Histograms showing the distribution of joke ratings and best-fitting beta distributions—obtained by maximum-likelihood estimation—for each of the 100 jokes (# indicates a joke’s number). The average rating ranges from *μ* = −3.43 to *μ* = +3.82, with a standard deviation of **σ** = 1.47 across those means. The average standard deviation is **σ** = 4.38 across jokes. Most of the jokes have unimodal distributions; only two clearly polarized the audience (jokes 71 and 75), as reflected by their bimodal distributions. The jokes are presented in decreasing order of their average rating (shown as *mu* in the panel titles). The x-axis shows the ratings and the y-axis shows normalized densities.

### 10.2 Comparisons between Strategies

#### 10.2.1 Comparing the Strategies Based on Individuals’ Taste Environments

Studies investigating different strategies’ performance across environments often pit the strategies against each other in informative match-ups [9]. To gain a more detailed understanding of how an individual’s learning environment influences the strategies’ performance, we compared pairs of strategies and studied the extent to which the strategies outperform each other for each of the 14,000 individuals.

Figure S3 shows scatterplots with all individuals positioned on axes of mean taste similarity with the crowd and dispersion of taste similarity with the crowd (as in Figure 3). However, rather than plotting the best-performing strategy for each individual as a function of experience, we now plot which of the two strategies in a comparison performed better for each individual. Each panel shows a different pair of strategies.

The first panel pits the performance of the *doppelgänger* strategy against the *whole crowd* strategy. For low levels of experience (left panel), the *whole crowd* strategy dominates for people whose preferences are positively correlated with the crowd, whereas the *doppelgänger* strategy performs better for individuals with alternative taste (i.e., negative mean taste similarity), where the *whole crowd* strategy performs worse than chance. As people become more experienced, the *doppelgänger* strategy starts to outperform the *whole crowd* strategy, also for people with more mainstream tastes. In particular, it has an edge for people who have higher dispersion in taste similarity with their peers. For these people, (i) it is more likely that similar individuals exist in the population and (ii) the strategy has better chances of identifying these very similar others.

A similar pattern of results can be observed in the second panel, which compares the *similarity-weighted crowd* strategy with the *similar crowd* strategy. The *similar crowd* strategy performs slightly better for individuals with high mean taste similarity with the crowd and low dispersion in taste similarity with their peers. The *similarity-weighted crowd* strategy performs notably better for individuals with high mean taste similarity and high dispersion in taste similarity. Note that the *similarity-weighted crowd* strategy is the only one that assigns negative weights to individuals perceived to have alternative taste (i.e., their preferences are negatively correlated with those of the social learner).

The third panel compares the *doppelgänger* strategy with the 10-person *clique* strategy. The *clique* strategy proves to be the better performing strategy almost across the board, essentially dominating the *doppelgänger* strategy. The only pocket of individuals for whom the *doppelgänger* strategy performs best are those with alternative tastes and low dispersion in taste similarity. This pocket reduces in size with experience.

The fourth comparison focuses on the 10-person *clique* strategy and the *similar crowd* strategy. For low levels of experience, the *similar crowd* strategy outperforms the *clique* strategy for almost the entire population, but as the level of experience increases, individuals with low average correlations with the crowd and high dispersion in taste similarity start to perform better when relying on their small *clique*.

Finally, we compared the *similarity-weighted crowd* strategy with the *whole crowd* strategy. For low levels of experience, the *whole crowd* strategy performs better for individuals with positive correlations with the crowd. As the level of experience increases, the *similarity-weighted crowd* strategy starts to outperform the *whole crowd* strategy for all individuals except for those with very low dispersion in taste similarity (i.e., those who benefit the least from distinguishing between more and less similar individuals).

**Figure S3:**
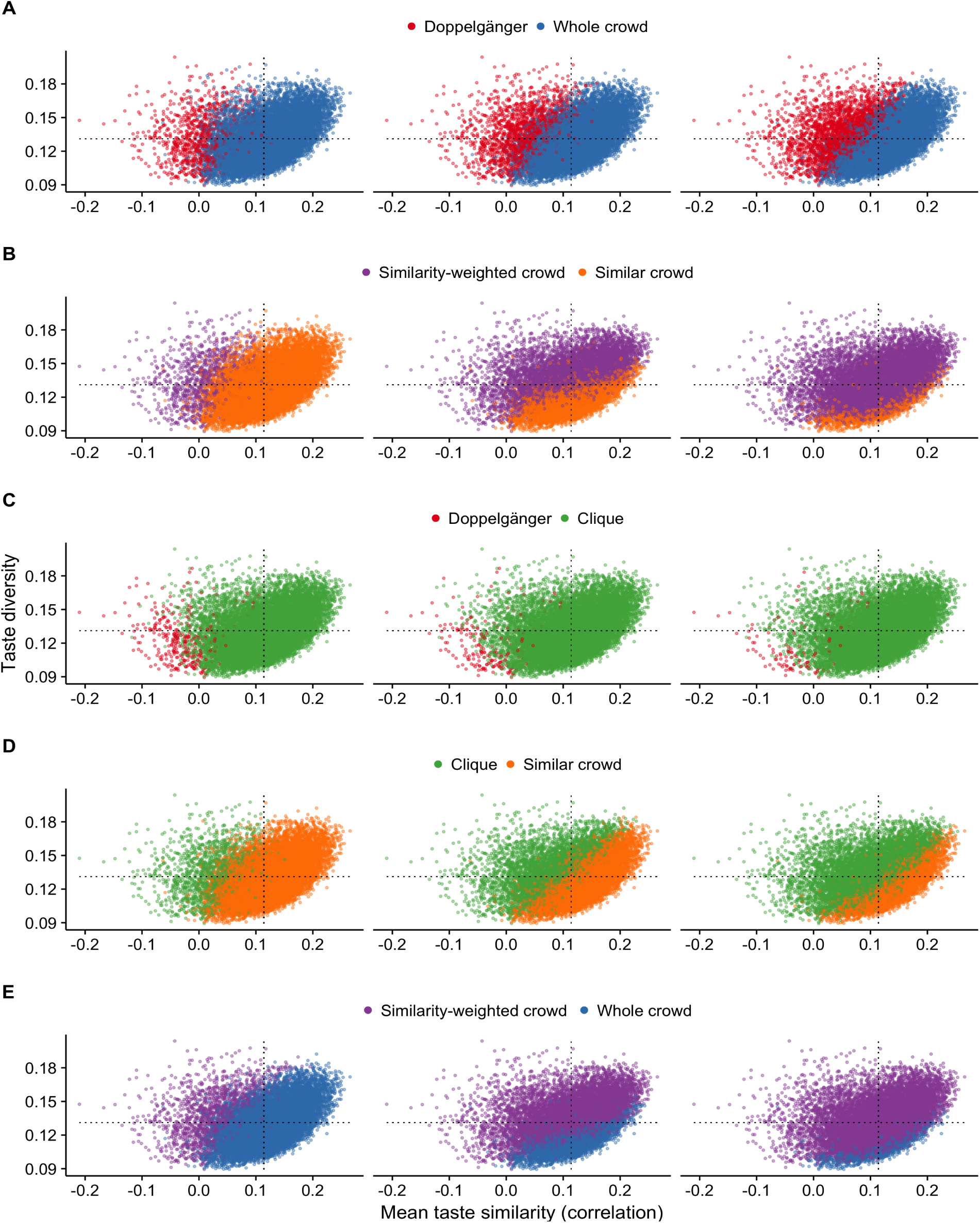
Comparing the strategies based on individuals’ taste environments. The figure is similar to Figure 3, but instead of plotting the best-performing strategy for each individual, it shows which of two strategies performed better for each individual. Each point represents one of the 14,000 participants. Each individual is positioned according to their mean taste similarity with all other 13,999 individuals (x-axis) and the dispersion in taste similarity with other individuals (i.e., the standard deviation of these correlations; y-axis). Each panel shows three experience levels from left to right: 10, 25, and 75 of options experienced. Panel **A:** *doppelgänger* versus *whole crowd*; Panel **B:** *similarity-weighted crowd* versus *similar crowd*; Panel **C:** *doppelgänger* versus *clique;* Panel **D:** *clique* versus *similar crowd*; Panel **E:** *similarity-weighted crowd* versus *whole crowd*. Results are based on averaging across 2,000 repetitions of the simulation. Standard errors for all individuals, strategies, and experience levels ranged between 0.00071 and 0.00287 with a median of 0.0028.

#### 10.2.2 Crossing Point Between Strategies as Experience Increase

**Figure S4:**
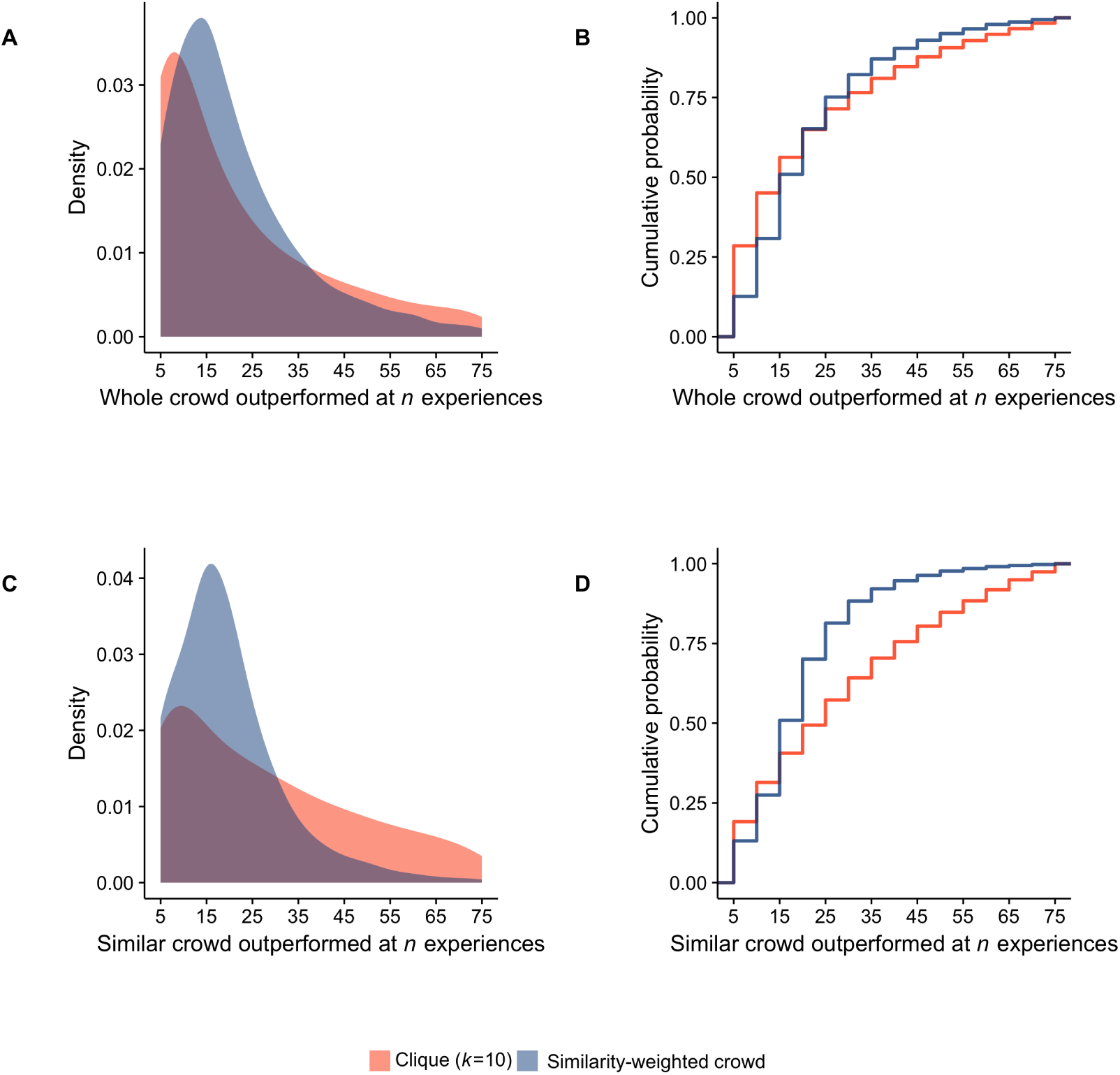
The proportion of people for whom the *clique* and the *similarity-weighted crowd* strategy outperformed the *whole crowd* strategy (panels A and B) and the *similar crowd* strategy (panels C and D) after a given number of experiences. Panels A and C show Gaussian kernel smoothed densities. Panels B and D show empirical cumulative distribution functions; the curves are step shaped because we varied the amount of experience in discrete steps (from 5 to 75 in steps of 5). Results are based on averaging across 2,000 repetitions of the simulation.

To better illustrate when the performances of strategies relying heavily versus loosely on similarity cross as experience increases, we focused on a few informative comparisons between pairs of strategies. Specifically, we calculated the proportion of people for whom two best-performing similarity-based strategies (i.e., *similarity-weighted crowd, clique*) surpassed the aggregation-heavy strategies (i.e., *whole crowd* and *similar crowd*) (Figure S4). The *similarity-weighted crowd* strategy outperforms both aggregation-heavy strategies for more than half of the population once 15 items have been experienced; the *clique* strategy outperforms the *whole crowd* strategy for more than half of the population once 20 items have been experienced and the *similar crowd* strategy once 25 items have been experienced.

### 10.3 When Similar Others Have Experienced Only Some of the Options

In the baseline simulation, we assumed that the decision makers’ peers had experienced all of the available items. This was a convenience assumption. In real life, a person’s friends and acquaintances have not experienced all items in a specific domain but only a limited number of them. Thus, people typically have to deal with a sparse matrix of peer experiences from which to estimate similarity and assess the value of future experiences. The same applies to most recommender system datasets: Individuals have experienced only small subsets of items that are curated by the system. A typical example is the MovieLens recommender system—one of the earliest recommender systems, developed and sustained by the GroupLens group at Minnesota University.^1^ It contains several million individual evaluations but only a small fraction of users have evaluated even the most popular items, making the dataset highly sparse. Likewise, the dataset released for the Netflix competition—although it contains more than a 100 million evaluations—is very sparse even for the most popular movies.

**Figure S5:**
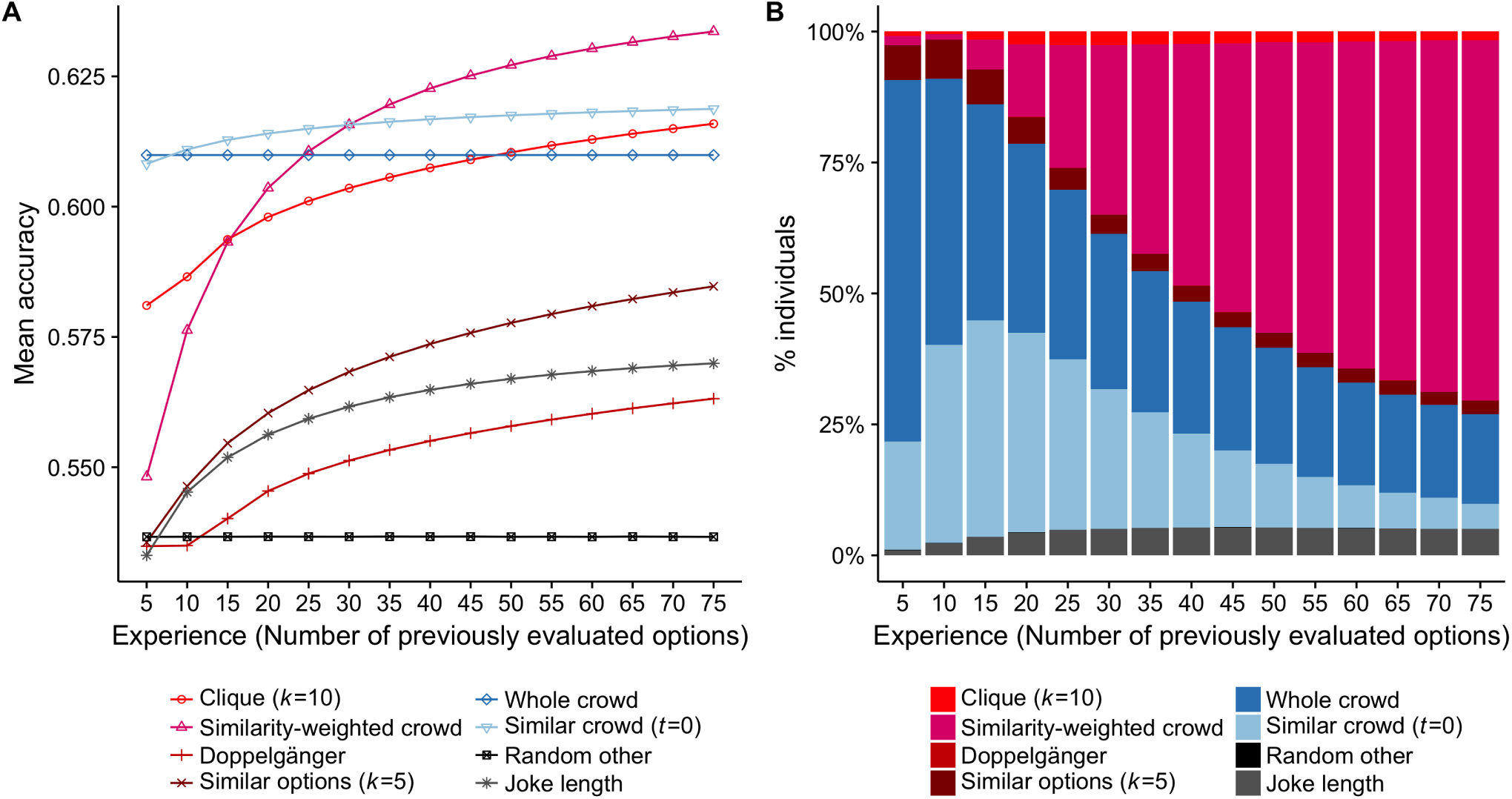
The effect of sparse matrices on the learning curves of the social learning strategies. This figure is structured identically to Figure 1 and displays the results of a simulation variant in which half of the items’ evaluations were deleted at random from the matrix before running the simulation. Relative to the original results (Figure 1), the crossing points between the similarity-heavy and the aggregation-heavy strategies have moved to the right. This result indicates that in many real-world, sparse domains, individuals need to experience even more options (than our baseline simulation would suggest) to effectively use strategies that rely heavily on similarity.

To investigate the sensitivity of our results to the assumption of a non-sparse matrix (as in our main simulation study), here we relax this assumption by randomly removing half of the values from the opinion matrix before each run of the simulation. There are at least two ways in which sparsity can reduce the performance of social learning strategies. First, people have fewer data available from which they can estimate similarity between themselves and their peers or items. For example, it could be that Sofia does not know how Bob has evaluated *Superman* and *Spiderman*, either because Bob has not watched the movies yet or because he has not talked about the movies to Sofia. Thus, Sofia will have fewer observations on the basis of which to estimate her taste similarity with Bob. This limitation will negatively affect all the strategies that rely on similarity estimation. Second, some (seemingly or truly) similar others might not have evaluated some of the items and will therefore not inform the prediction of how much the target individual will enjoy an item. For strategies that rely on a fixed number of similar others (e.g., *clique*) or items, we assumed that the missing value of the similar individual (or item) is substituted by the next most similar individual (or item) from the crowd (or collection of similar items). In the case of the *doppelgänger* strategy, Bob will be substituted by John, who is the next most similar individual. Besides these differences in strategy implementation, the details of the simulation remained unchanged.

Figure S5A shows that (relative to the main simulation; Figure 1) the strategies that rely less on similarity perform best more often for even more experienced individuals. That is, the crossing points at which, on average, similarity-heavy strategies overtake aggregation-heavy strategies have moved further to the right. The *similarity-weighted crowd* strategy outperforms the *whole crowd* strategy after 30 experiences and the *similar crowd* strategy after 45 experiences. The *clique* strategy outperforms the *whole crowd* strategy after 50 experiences and it is still outperformed by the *similar crowd* strategy after 75 experiences. The other two strategies that rely on similarity are affected in a similar way. The *doppelgänger* strategy now performs consistently worse than the *joke-length* strategy, while the *similar options* strategy performs, on average, well below the *whole crowd* strategy. Note that the performances of the *whole crowd* and *similar crowd* strategies remain almost completely unaffected in terms of absolute performance by the sparsity introduced in this simulation. The only difference is that they now use less data to calculate the average qualities of objects. These results are also reflected in the number of individuals for whom a certain strategy performed best. As seen in Figure S5B, the aggregation-heavy strategies perform best for a much larger fraction of individuals. The *similarity-weighted crowd* strategy becomes the best-performing one only after 40 options have been experienced.

### 10.4 Varying the Size of the Community

#### 10.4.1 Accessing Small Communities of Close Friends

**Figure S6:**
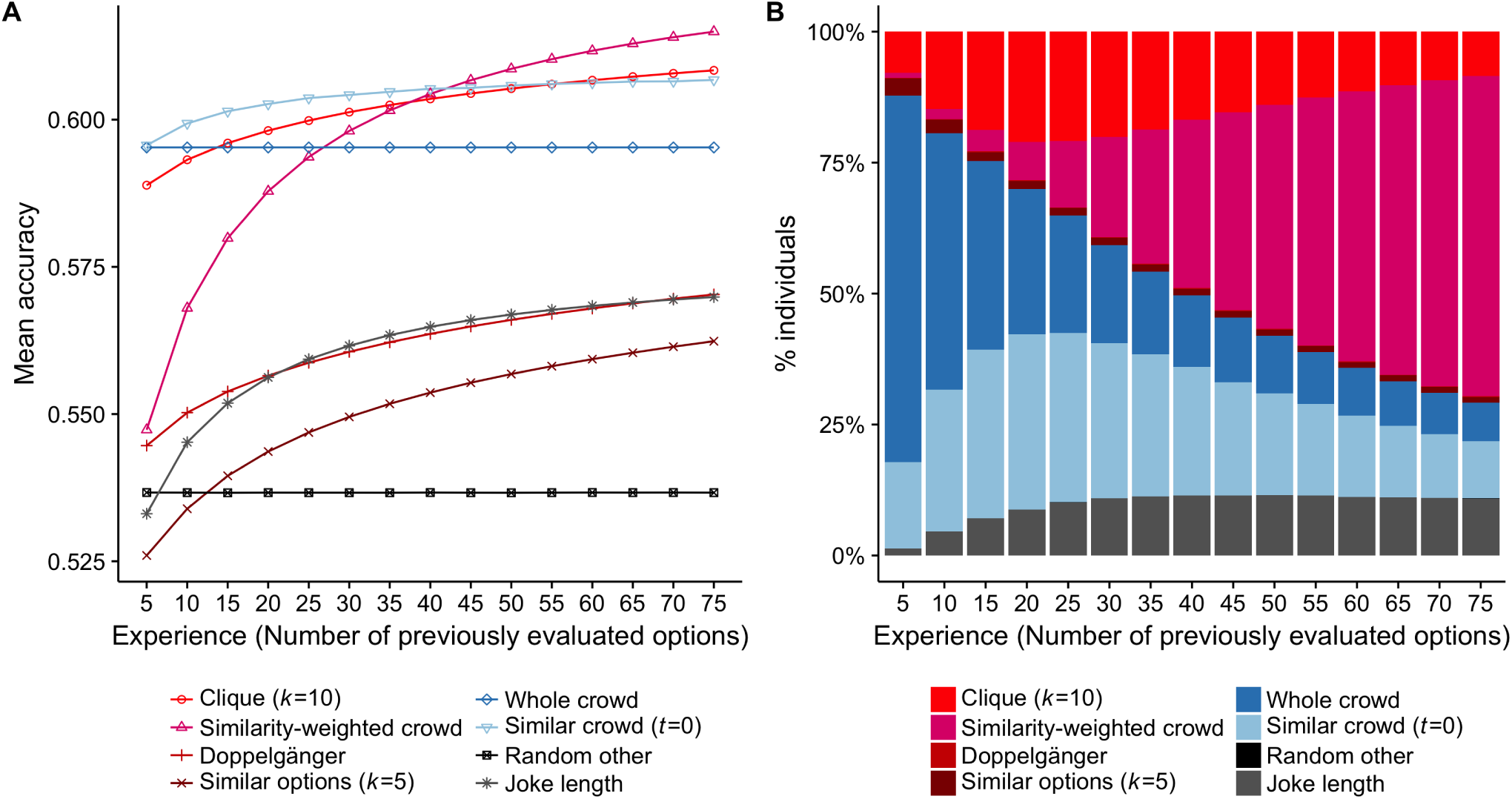
Performance of the social learning strategies when decision makers have access to only a small community of 25 individuals. This figure is structured identically to Figure 1.

So far, we have assumed that people have access to a community of only 249 other individuals, an assumption that reflects both cognitive and social constraints in everyday life [3]. One could argue, however, that this assumption is rather generous, as in many cases it will be costly to ask friends and acquaintances about their experiences. To account for this possibility, we restricted the number of individuals in a community to 25 (24 plus the target individual) and reran the simulation, keeping the rest of the simulation setup exactly as in the baseline simulation study reported in the main text. When the size of the community is reduced, the performance of all strategies declines, but the key results and conclusions of the simulation remain the same. The main change is that the crossing points at which, on average, similarity-heavy strategies overtake aggregation-heavy strategies have moved further to the right. The *similarity-weighted crowd* strategy outperforms the *whole crowd* strategy after 30 experiences and the *similar crowd* strategy after 45 experiences. On the one hand, the performance of the aggregation-heavy strategies drops by about 1 percentage point uniformly across the levels of experience. On the other hand, the performance of the *similarity-weighted crowd* strategy now increases faster as a function of experience, presumably because the correct estimation of similarities is more crucial in a crowd of 24—the tastes of different individuals do not cancel each other out as easily. The *clique* strategy now outperforms the *whole crowd* strategy after 15 experiences and it needs 45 experiences to outperform the *similar crowd* strategy. The *similar options* strategy is most strongly affected by the smaller size of the community, as the amount of information available for the strategy to evaluate similarity between items is now drastically reduced.

#### 10.4.2 Accessing Large Communities: The Whole Population as the Pool of Information

**Figure S7:**
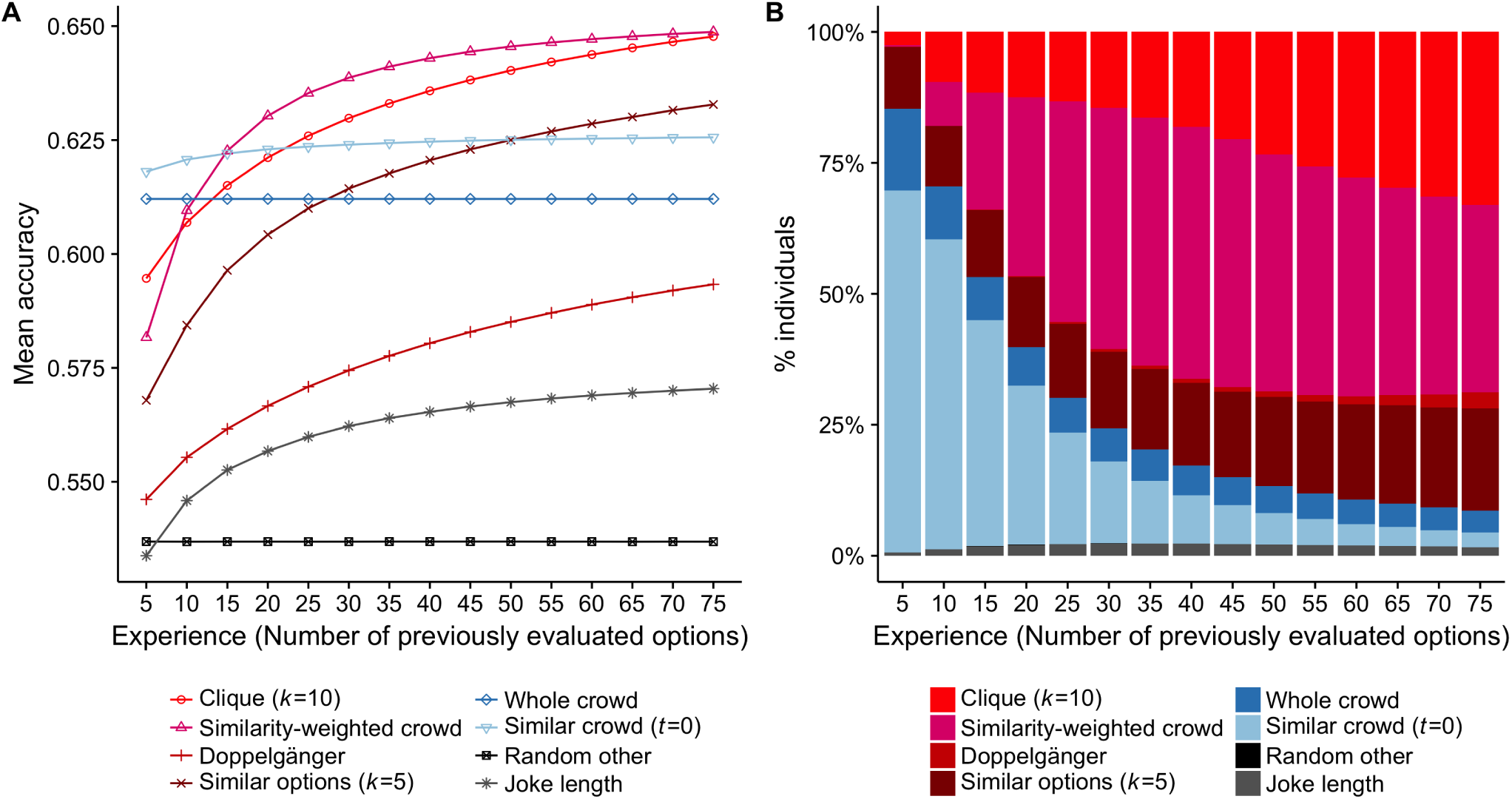
Performance of the social learning strategies when decision makers have access to all 14,000 individuals in the matrix. This figure is structured identically to Figure 1. Results are based on averaging across 2,000 repetitions of the simulation.

In most operational recommender systems, the collaborative filtering algorithms seek similarity patterns among several thousand users. Would having access to a much larger number of similar others substantially change the results presented so far? The answer is no (Figure S7). The performance of all strategies improves by a few percentage points, but the ranking of the strategies according to their performance remains almost the same. The main difference from the baseline simulation is that the number of experiences required for similarity-heavy strategies to outperform aggregation-heavy strategies is somewhat lower (i.e., the crossing points are shifted slightly to the left). Ten to 15 experiences are required for the *similarity-weighted crowd* strategy to outperform the *whole crowd* and *similar crowd* strategies. This result also illustrates the fundamental nature of the cold-start problem in recommender systems [4]. Even in domains where millions of other people have evaluated the options, at least some initial experiences are required in order to make meaningful predictions about any target user. Accurately estimating similarities between users will always be challenging when the estimates are based on only a small number of common experiences, even in cases where similarity patterns can be sought among a very large number of users.

### 10.5 Considering the Opinion of Less Similar People Can Improve Predictions

**Figure S8:**
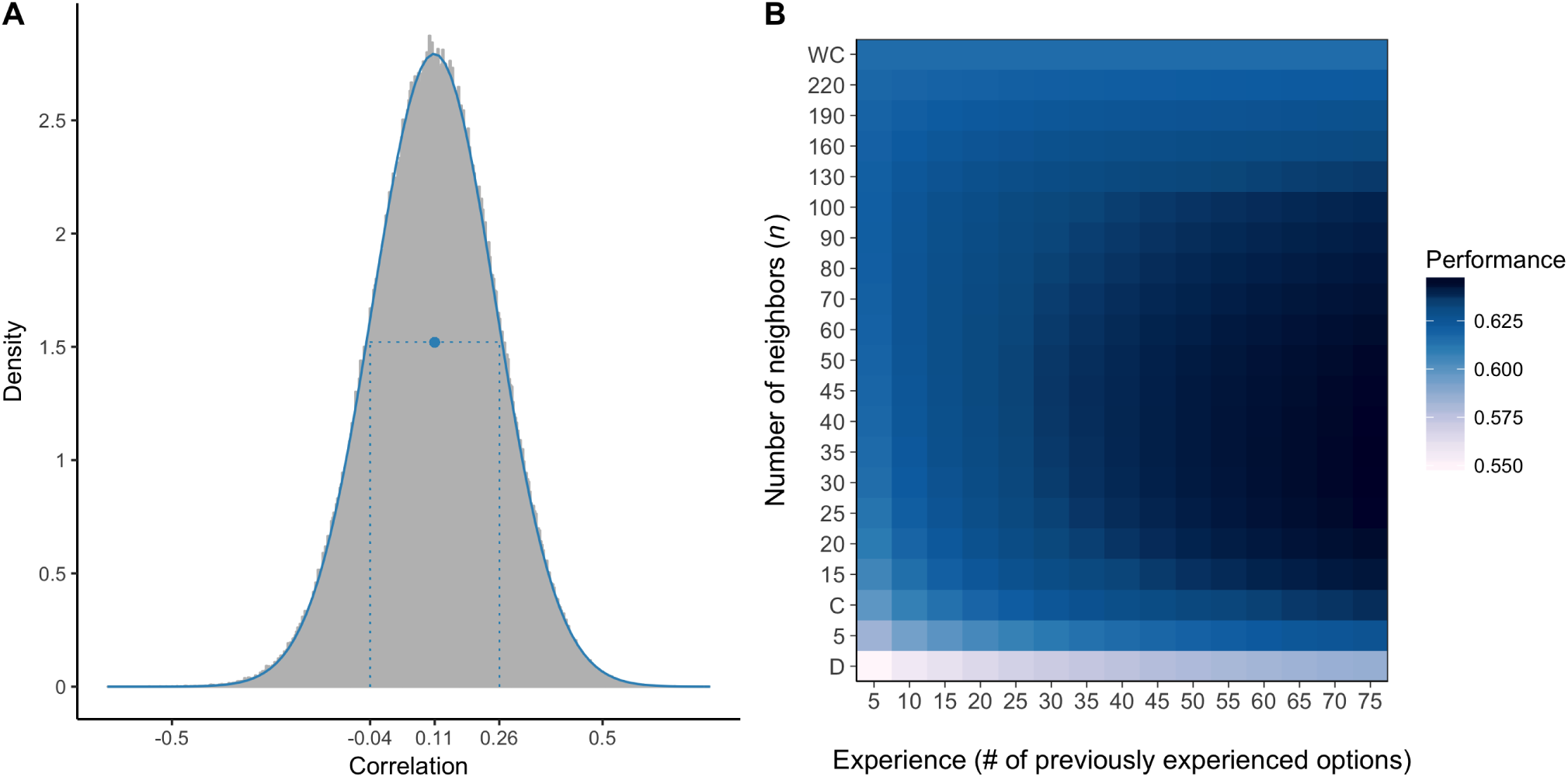
**Panel A:** The distribution of correlations among a random sample of 1,000 individuals (999,000 correlations in total). The dot shows the mean correlation and the two vertical lines indicate one standard deviation distance from the mean. **Panel B:** Performance of the *clique* strategy (i.e., aggregating the tastes of the *k* most similar individuals in the community) for different sizes of the clique (*k*). The top row of the matrix (WC) corresponds to the *whole crowd* strategy, the bottom row to the *doppelgänger* strategy (D), and the third row from the bottom to the *clique* (C) strategy as described in the main text (with *k* fixed at 10). The optimal number of similar others decreases with experience from 100 to about 30.

People who have not yet acquired any experience are necessarily unaware of their similarity with other individuals. People following the choices of a random individual should perform better than chance because, by statistical necessity, a majority of the population is expected to have a positive mean taste correlation. The extent to which the *random other* strategy will outperform chance depends on the mean correlation of tastes among different individuals (see Figure S8A). Decision makers’ best bet in that case is to use the *whole crowd* strategy to predict which of two options they will prefer. As they acquire some experience, they can use it to calculate their similarity with their peers. For decision makers with limited experience, relying on only the most similar other through the *doppelgänger* strategy marginally outperforms the *random other* strategy. Taking into account additional—although less similar—people and averaging their recommendations markedly improves performance (Figure S8). When using the *clique* strategy, the size of *k* (i.e., the number of neighbors whose evaluations are averaged) can be seen as a *hyperparameter* that needs to be chosen beforehand (in Figure 1A we fixed *k* at 10). With as few as five experiences, people can reduce the number of individuals they rely on from 249, which corresponds to the whole crowd, to 100. Once all 75 options have been experienced, performance peaks at moderately sized cliques (approximately 30).

### 10.6 Learning Curves for Six Prototypical Individuals

**Figure S9:**
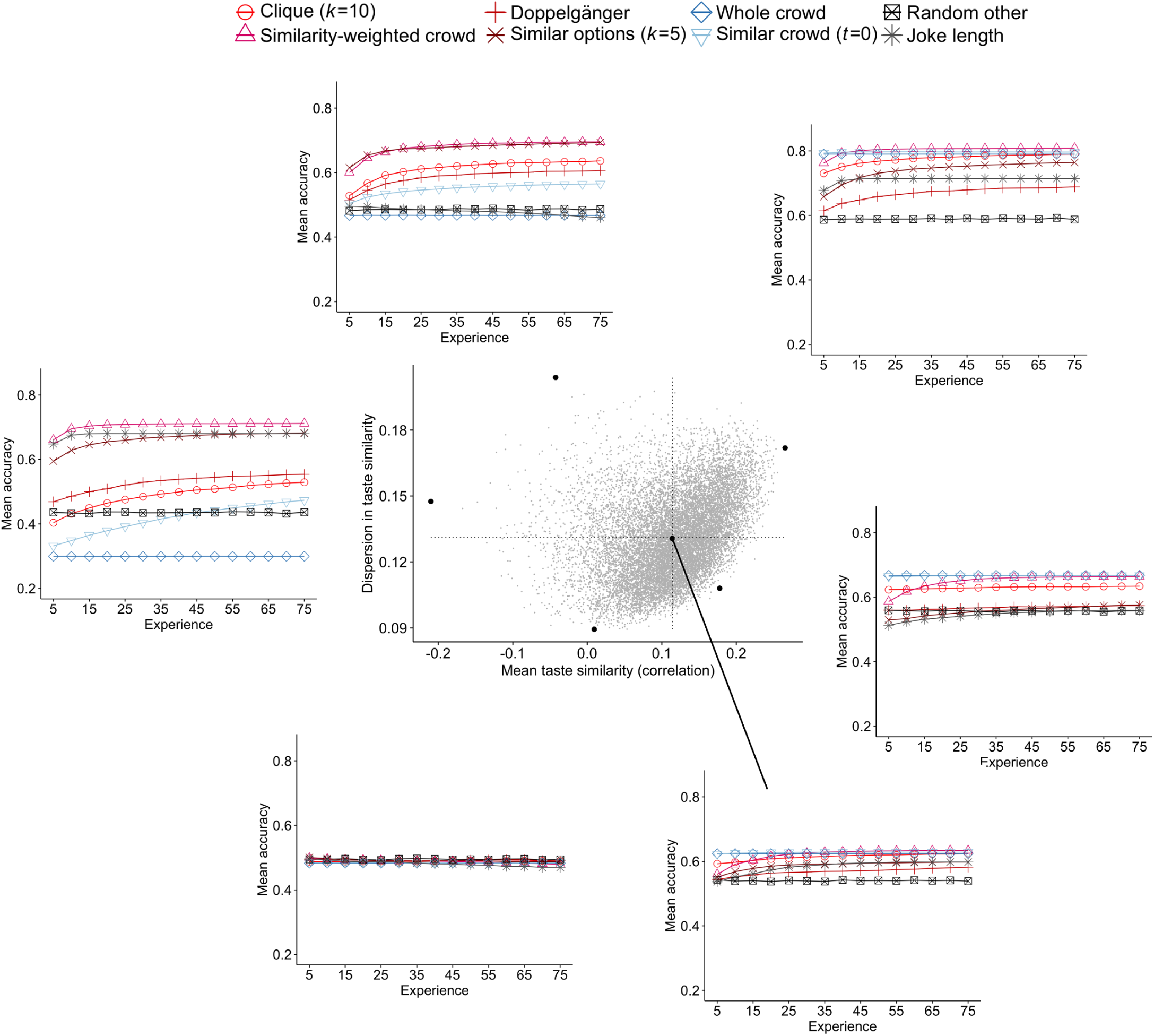
The performance of the learning strategies for six prototypical individuals selected from different regions of the two-dimensional space defined by mean taste similarity and dispersion in taste similarity. The scatterplot (at the center of the figure) shows all 14,000 individuals; the black dots represent the six prototypical individuals. Each individual is positioned according to their mean taste similarity with all other 13,999 individuals (x-axis) and dispersion in taste similarity with other individuals (i.e., the standard deviation of these correlations; y-axis). The six outer graphs illustrate the performance of the social learning strategies for the six prototypical individuals as their experience increases (i.e., with the number of jokes experienced and evaluated); see, for example, Figure 1A for an explanation of the strategy legend. The learning curves of individuals drawn nearby prototypical others were very similar to the learning curves of those prototypes. Results are based on averaging across 2,000 repetitions of the simulation. Standard errors for all individuals, strategies, and experience levels ranged between 0.00071 and 0.00287 with a median of 0.0028.

As illustrated by Figures 2 and 3, the performance of different strategies and their potential for improvement with increasing levels of experience varies substantially across different people in the population and depends on how mainstream or alternative an individual’s tastes are, as well as on the level of dispersion in taste similarity with the other people in the group.

To better connect the aggregate-level analyses with the individual-level analyses, we sampled six prototypical individuals from different regions of the two-dimensional space defined by mean taste similarity and dispersion in taste similarity and investigated how the different social learning strategies performed for these people as a function of experience. We used the following sampling process: First, we selected individuals from the four extremes of the space (highest and lowest mean taste similarity, and highest and lowest dispersion in taste similarity). We also sampled the individual who was located closest to the mean values of mean taste similarity and dispersion in taste similarity in the population, representing the central person. This selection procedure left the lower-right quadrant of the space (high mean taste similarity and low dispersion) unoccupied, so we selected an additional individual from the outermost edge of this population cluster. For these selected individuals, the strategies differ greatly in their performance and potential for improvement when more experiences are used to estimate similarity. For example, all strategies perform at chance level for a very idiosyncratic individual (bottom-left graph), whereas several strategies correctly predict about four-fifths of the pair comparisons for the individual with the highest mean taste similarity in the population (upper-right graph).

Similarity-heavy strategies, such as the *clique* and *similarity-weighted crowd* strategies, improve more as a function of experience for individuals with alternative tastes and higher dispersion in taste similarities. Although the *whole crowd* and *similar crowd* strategies perform worse than chance level for individuals with alternative tastes (i.e., negative mean taste similarity), similarity-heavy strategies perform remarkably well for those individuals, even after just a few experiences, and performance improves further as experience increases. In contrast, for people with high mean taste similarity and low dispersion in taste similarity, there is less room for improvement in the estimates of similarity (i.e., compare the central individual and the mainstream individual with low dispersion in taste similarity with individuals with higher dispersion in taste similarities). Aggregation-heavy strategies, such as the *whole crowd* and *similar crowd* strategies, already perform relatively well for these people and for some individuals they are not outperformed at any level of experience by any other strategy (e.g., individuals on the edge of the lower right quadrant).

Figures 2 and S10 show that for individuals located in nearby coordinates in the two-dimensional space (mean taste similarity vs. dispersion in taste similarity), the performance of each of the strategies is very similar (across the three levels of experience depicted: 10, 25, and 75 options). Furthermore, Figure 3 reveals that for three different level of experiences the same strategies tend to perform best for individuals drawn from nearby coordinates in our space. Thus, we are confident that the prototypical individuals reflect strategy performance and learning patterns for other people sampled from nearby coordinates in the two-dimensional space we defined. To verify this observation, we also sampled several people from the regions in which the six prototypical individuals were drawn and examined the learning curves. Indeed, the social learning strategies for individuals selected from nearby coordinates in the two-dimensional space had very similar performance levels at different levels of experience.

### 10.7 Error of the Strategies for Different Individuals and Levels of Experience

As seen in Figure 1, the performance of all social learning strategies—except the *whole crowd* and the *random other* strategies—improves with experience. Furthermore, from Figure 2 we see that performance varies considerably across individuals, depending on their mean taste similarity with others and the dispersion of similarities between an individual and her peers. Does acquiring more experience lead to an equal improvement in performance across individuals? Figure S10 plots the performance of each strategy and individual at three levels of experience (10, 25, and 75 experiences, respectively). For all strategies—except the *whole crowd*—there is much more learning potential (i) for individuals with less mainstream or even alternative tastes (i.e., those with negative mean taste correlations) and (ii) for people whose taste similarities with their peers are more dispersed. For all strategies, the largest improvements occur in the upper left quadrants of Figure S10’s panels, followed by the upper right quadrants, demonstrating that the people’s dispersion in taste similarities is crucial for learning. Clearly, in a more diverse peer group there are more possibilities to identify very similar individuals. In contrast, the lowest levels of learning occur for individuals who have low dispersion of similarities and a very low average similarity with their peers (bottom part of the graph in the lower left quadrant.)

### 10.8 Bias–Variance Decomposition

#### 10.8.1 The Bias–Variance Dilemma

The bias–variance decomposition was popularized in machine learning in a seminal paper by Geman, Bienenstock, and Dursat [7]. Yet the first explicit implementation of a bias–variance analysis that we are aware of took place at least two decades earlier, when Hoerl and Kennard [8] used such an analysis to develop *ridge regression*, a biased yet robust alternative to classic multi-linear regression models that rely on ordinary least squares (OLS) methods. The starting point for a bias–variance analysis is to decompose the error into three components:

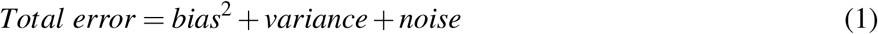

*Bias*^2^ represents the difference between the average prediction of the model and the underlying truth. As Hoerl and Kennard illustrated in their pioneering work, a model or an estimator can be provably unbiased and still suffer from higher total error than a more biased estimator. The reason is that the unbiased estimator may have a larger variance, that is, the predictions of the algorithm are more sensitive to the training data that has been sampled. Finally, noise represents the irreducible error in the function generating the data and delineates the ultimate performance limit for any model. Estimating the irreducible error in real data is challenging because the function generating the data is usually not known. However, because the variance component is defined as the dispersion around the average prediction of the model, the variance component can be estimated without knowing the function generating the data. Subtracting the variance from the total error then results in a mixed term reflecting *bias*^2^ plus noise. Therefore, and as often done in the literature, we will decompose the total error into the variance component and this mixed term reflecting *bias*^2^ plus noise.

To further clarify the bias and variance components of the error, let us consider the dartboard illustrations in Figure S11. Ideally, we would like our model to predict the two-dimensional coordinates corresponding to the middle of the target (the exact middle of the dartboard). Let us assume that the blue dots on the boards represent individual predictions of our model based on different training samples. The model used on the upper right panel is almost unbiased. The average prediction is close to the middle of the target. Yet, the individual predictions deviate markedly from one another and also from the center of the target. The observed variance of the individual predictions leads to a relatively high total error. Ideally, we would like our model to have both low bias and low variance, as illustrated by the upper left dartboard. Yet, in many real-world prediction problems, this aspiration is unattainable.

**Figure S10:**
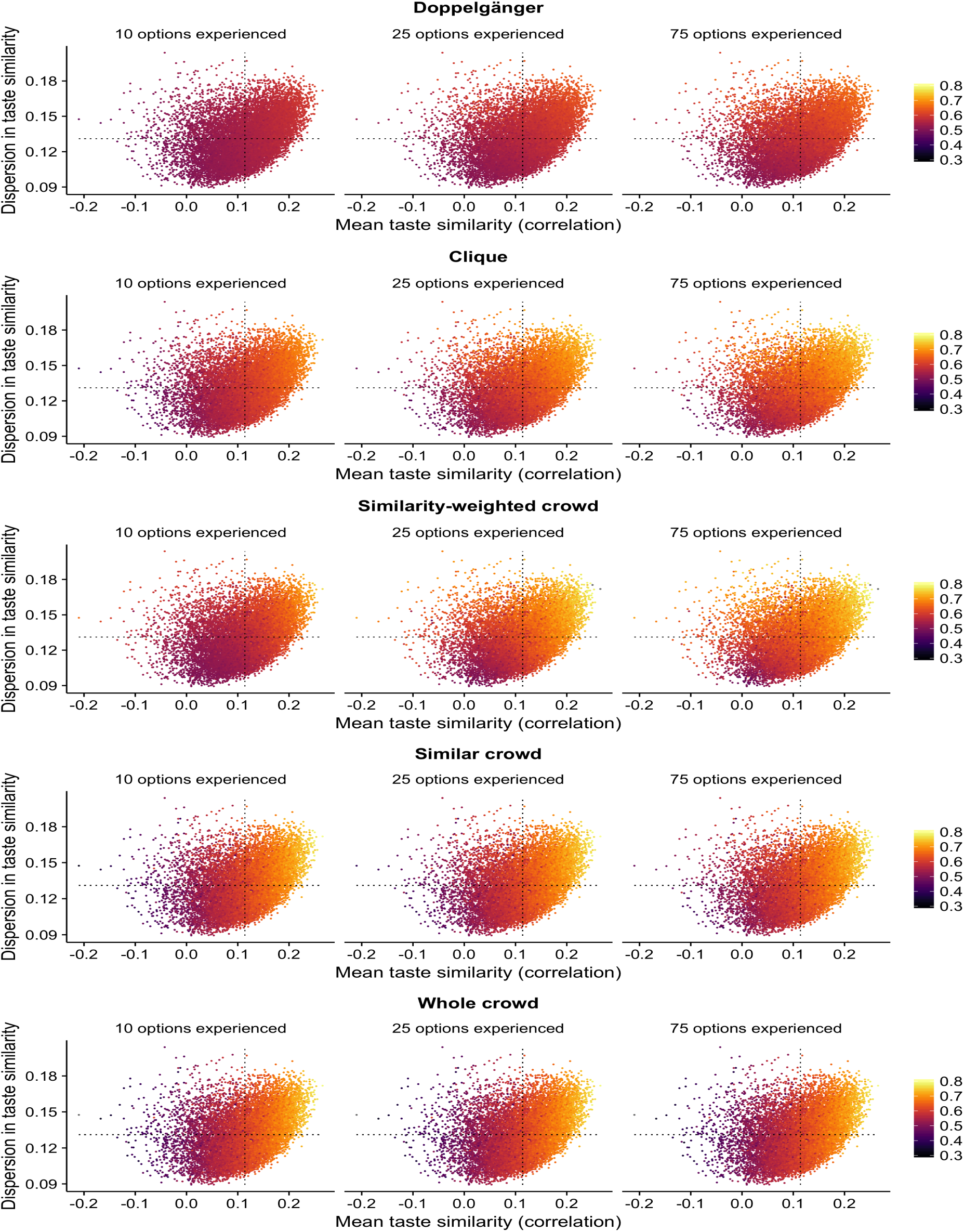
Strategies’ performance after experiencing 10, 25, and 75 options, respectively, depending on how people’s tastes correlate with those of others in the population. The scatter plots represent the strategies’ performance for each of the 14,000 individuals (i.e., proportion of correct predictions, color coded). Each point represents one of the 14,000 individuals and thus represents a unique prediction environment for social learning strategies for matters of taste. Each individual is positioned according to their mean taste similarity with all other 13,999 individuals (x-axis) and the dispersion in taste similarity with other individuals (i.e., standard deviation of these correlations; y-axis). Dashed horizontal and vertical lines show the overall average of the mean correlations (0.11) and standard deviations (0.13). Results are based on averaging across 2,000 repetitions of the simulation.

The art of developing robust models for accurate out-of-sample predictions in real-world problems requires striking a good balance between bias and variance. Typically, very complex models with many free parameters are flexible and can often exactly fit the data that was used to train them. As the training data varies from one training sample to another, so do the resulting predictions of the model, leading to a high variance component. Simpler models tend to be more biased and less flexible (see the lower left target in Figure S11). Models that make exactly the same prediction regardless of the training set have no variance whatsoever but are likely severely biased (see the graph on the right of Figure S11, at the very left end of the spectrum). Very complex models, on the other hand, generally have very low bias. Their average predictions hit close to the target. Yet they suffer from a lot of variance (see the graph on the right of Figure S11, at the very right end of the spectrum).

**Figure S11:**
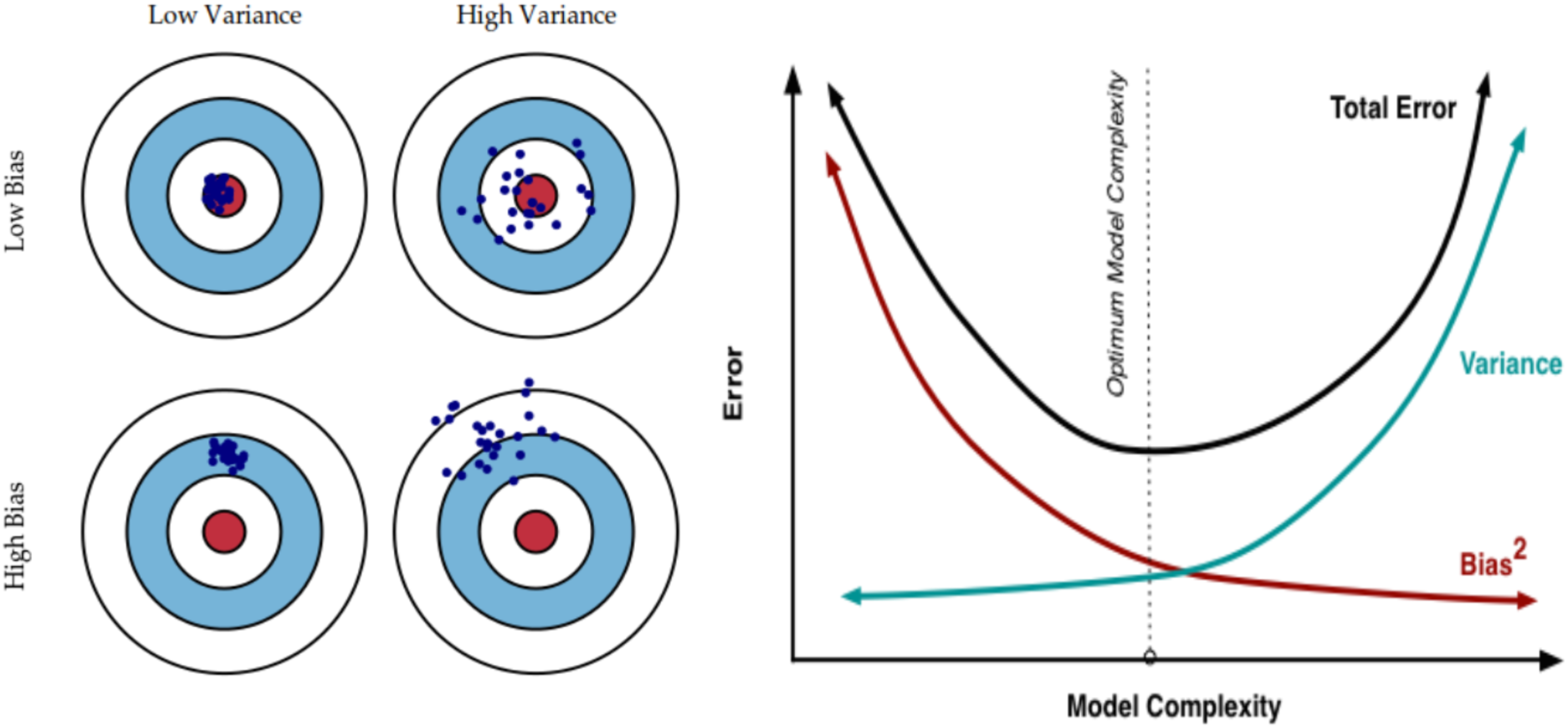
**Left**: An illustration of the bias–variance decomposition for continuous loss functions. A prediction method might be unbiased but still have a relatively high error (upper right dartboard) due to high variance, that is, the sensitivity of the method to the sample that was used to train the model. **Right:** As model complexity increases, the model becomes more flexible, allowing for lower bias. Yet more complex models often imply higher variance. These two illustrations were designed by Fortmann-Roe [5].

#### 10.8.2 Bias–Variance Decomposition for the Zero-One Loss Function

In this section, we illustrate how to decompose the error of a strategy into a bias and a variance component for binary categorization tasks. Whenever strategies are evaluated on pair comparisons (“Is it A or B?”), their decisions are either correct or incorrect. This evaluation function is also called the *zero-one loss function*, where correct decisions incur no loss (*L* = 0) and incorrect decisions incur a unit loss (*L* = 1). The average loss (or cost) of a prediction method *E* (*C*) is commonly referred to as the misclassification error. Although for continuous estimation tasks (often called “regression” problems in machine learning) there has been a consensus on how to implement the bias–variance decomposition (see also the previous section), this is not the case for binary categorization tasks [8, 10]. There have been several suggestions about the best way to implement the bias–variance decomposition for the zero-one loss function [12, 1,11, 14, 6, 10]. For an extensive study of the bias–variance decomposition on real data including problems with a zero-one loss function, please refer to ongoing work by Şimşek and Buckmann [13].

Here we adopt the method advanced by Kohavi and Wolpert [11], which is the most widely implemented method in the literature on categorization. All other decompositions have the drawback that the variance might be negative, and might not equal 0 for algorithms like the *whole crowd* strategy presented in this paper, which always make the same predictions irrespective of the particular set of training data. We consider this a significant advantage, as the resulting decomposition is intuitive and easily interpretable.

Kohavi and Wolpert [11] formalized the cost function for zero-one prediction problems and, with a few algebraic transformations, showed how to decompose the zero-one loss function into its components (i) bias, (ii) variance, and (iii) irreducible error. For a target function *f* and a training set of size *m*, the expected misclassification rate *E* (*C*) can be written as

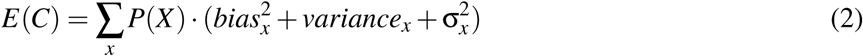

where

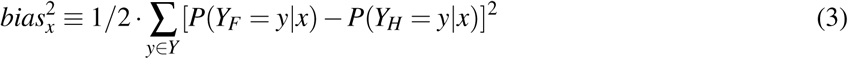

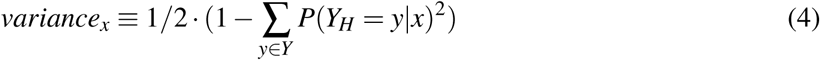

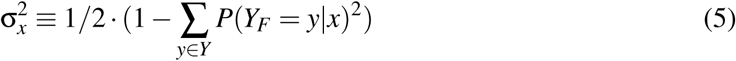

In the above equations, *X* and *Y* are the input and output spaces, that is, the space of possible feature values and categories or outcomes. *x* and *y* are particular instances in these spaces, that is, specific combinations of feature values and categories that can be used to either train or evaluate the model. The target function *f* is a conditional probability distribution *P*(*Y_F_* = *y*|*x*). This expression states that for feature values *x*, there is a given probability that the item will belong to category *y*. The hypothesis or model *h* generated by the learner is also a distribution *P*(*Y_H_* = *y*|*x*). This expression states the probability that the model constructed will predict a certain category *y* in the output space when it sees a feature vector *x*.

#### 10.8.3 Illustrating the Kohavi-Wolpert Bias–Variance Decomposition: Coin Flip Examples

Let us now illustrate how the above equations work when we predict whether the outcome of tossing a coin (fair or rigged) will be *heads* or *tails*. First, let us assume that we are dealing with a fair coin. There are two possible outcomes in *Y*, *heads* and *tails*. Then, let us assume that regardless of any information *x* that we might possess, it is impossible to predict at better than chance level.^2^

If we always predict tails regardless of the side information available *x*, the expected misclassification error (loss) will be equal to 0.5 (i.e., a fair coin will land heads in 50% of cases and therefore we will be wrong in 50% of cases). For either heads or tails, the *P*(*Y_F_* = *y*|*x*) = 0.5. Our hypothesis *P*(*Y_H_* = *y*|*x*), however, is that the coin is rigged and it always comes out tails (these are the two possible outcomes in space *Y*, unless the coin can stand on its edge). We may have arrived at that belief by falsely interpreting the available side information *x* (i.e., misreading an omen). For tails *P*(*Y_H_* = *y_tails_*|*x*) = 1 and for heads *P*(*Y_H_* = *y_heads_*|*x*) = 0. Thus, for both cases, we would be equally off target in our predictions for either of these hypotheses because [*P*(*Y_F_* = *y_tails_*|*x*) — *P*(*Y_H_* = *y_tails_*|*x*)]^2^ = [*P*(*Y_F_* = *y_heads_*|*x*) — *P*(*Y_H_* = *y_heads_*|*x*)]^2^ = 0.25. The *bias*^2^ term in the Kohavi–Wolpert decomposition measures the sum of the squared differences between the actual outcome and the algorithm’s average hypothesis, or predicted outcome. Following equation 3, by averaging the results from the two possible outcomes, we arrive at the contribution of *bias*^2^ to the overall error.

The variance captures the variability in the predictions generated by the algorithm. In this case, because the algorithm always predicts tails regardless of the information at hand, the variance of the model will be equal to 0. The irreducible noise σ^2^ corresponds to the error that even a Laplacian demon with perfect knowledge of the universe or the creator of an experiment, with precise knowledge of its setup, cannot avoid given the information available. Here, the noise is the randomness inherent in a coin flip. The σ^2^ is independent of the learning algorithm and captures the variance of the target function. In this scenario, it also equals 0.25 (0.5^2^). Now let us assume that the algorithm randomizes between heads and tails, assigning equal probabilities to both outcomes. Our model’s *bias*^2^ will equal 0. The classification error will be distributed equally between *variance* = 0.25 and σ^2^ = 0.25. Finally, let us assume that the coin is rigged and always comes up heads. If the decision maker randomizes her predictions between *tails* and *heads*, the misclassification error will again equal 0.5. It will now be divided equally between *bias*^2^ = 0.25 and *variance* = 0.25. If she always mistakenly predicts tails although the rigged coin always comes up heads, the misclassification error will be 1 and it will consist exclusively of *bias*^2^.

In empirical settings, the variance term measures the sensitivity of the algorithm to changes in the training data. To illustrate this point, let us assume that the coin is rigged and that it comes up heads in 0.65 of throws. Let us also assume that the decision maker has access to a sample of some past observations from our coin that she can use to forecast future coin tosses. Finally, let us assume that she has opted to use maximum likelihood estimation (MLE) to generate predictions about future outcomes. If her maximum likelihood estimate suggests that the coin is rigged to come up heads more often than tails (MLE for heads above 0.5), she will always predict heads; if it is below 0.5, she will always predict tails, while she would randomize when the MLE suggests that the coin is fair (MLE for heads is 0.5). Let us now assume that the decision maker has access to a sample of 10 observations from our coin. Her MLE approach would lead her to call heads if she saw more than 5 heads, tails if she saw less than 5, while she would randomize when she saw exactly five. We can simulate this decision process repeatedly (or use the expectation of the binomial distribution) to find out that our protagonist will predict heads 82.6 % of the times. The average error in that case is 0.394 (slightly larger than it would have been if she had always predicted heads). The variance accounts for 0.142, the irreducible error for 0.228, and the *bias^2^* for 0.025. If she had access to 20 observations, she would predict heads 91 % of the time and variance would account for 0.082 of the overall error, which is now 0.378. If, in contrast, she had access to only 3 observations, she would predict heads only 73% of the time and the variance would account for 0.197 of a total error of 0.43.

#### 10.8.4 Calculating Bias and Variance from Data

In the previous section, we illustrated how the bias–variance decomposition works in a scenario where the underlying truth is known and the strategies do not have any free parameters (but there is inherent stochasticity in some of their predictions). We now show how to apply the above concepts when using different training samples *d*. First, let us assume that there is a target function *f* that governs the statistical relations in the environment and from where the samples *d* are drawn, and let us define the size of these training samples as *m*.

As we have already seen, *P*(*Y_F_* = *y*|*x*) indicates that the probability that the function takes the value *y* for a feature vector *x*. Using the concepts introduced above, we can rewrite *P*(*Y_F_* = *y*|*x*) as *P*(*Y_F_* = *y*|*f,m,x*), which in turn can be written as 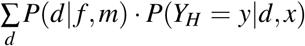. The term *P*(*d*|*f,m*) in the last expression represents the probability of generating the training data *d* from target function *f*, and *P*(*Y_H_* = *y*|*d,x*) represents the probability that the learning algorithm will make prediction *y* after seeing the feature vector *x* when trained with training sample *d* of size *m*. Therefore, *P*(*Y_H_* = *y*|*x*) is the average *Y* value over all training sets that is predicted when the learner is exposed to a feature vector with values *x*.

In most real-life problems, decision makers do not know the underlying function *f*. They only have access to limited number of *y* and *x* pairs in a dataset. Following Kohavi and Wolpert [11], we define a training space *D* from where different samples *d* will be drawn and an evaluation space *E*, representing the instances that are kept apart in order to evaluate the model. Each draw from *D* is equiprobable, thus we can simply average the predictions for different training samples for each instance with feature profile *x*. We can calculate the overall error and the bias and variance components by going back to equation 2 and averaging over the error, bias, and variance terms of each individual instance in the evaluation space *E*.

#### 10.8.5 Simulation Procedure for the Empirical Bias–Variance Decomposition

The Jester dataset has 100 items in total. Given that some of the items need to be reserved for the test set (for cross-validation), there is a limit to how large the set of training items in the bias–variance decomposition simulation can be. Ideally, the number of training examples should be small in relation to the entire training space (i.e., the pool of jokes from which training examples are drawn). Otherwise, the training samples might contain many similar objects, leading to an underestimation of the variance component. To avoid this issue, we opted to study the decomposition at only one level of experience (i.e., number of previously experienced items).^3^ To study how the error decomposes into bias and variance for each individual in the Jester dataset, we replicated the procedure put forward by Kohavi and Wolpert [11], with a few small modifications:

**Figure S12:**
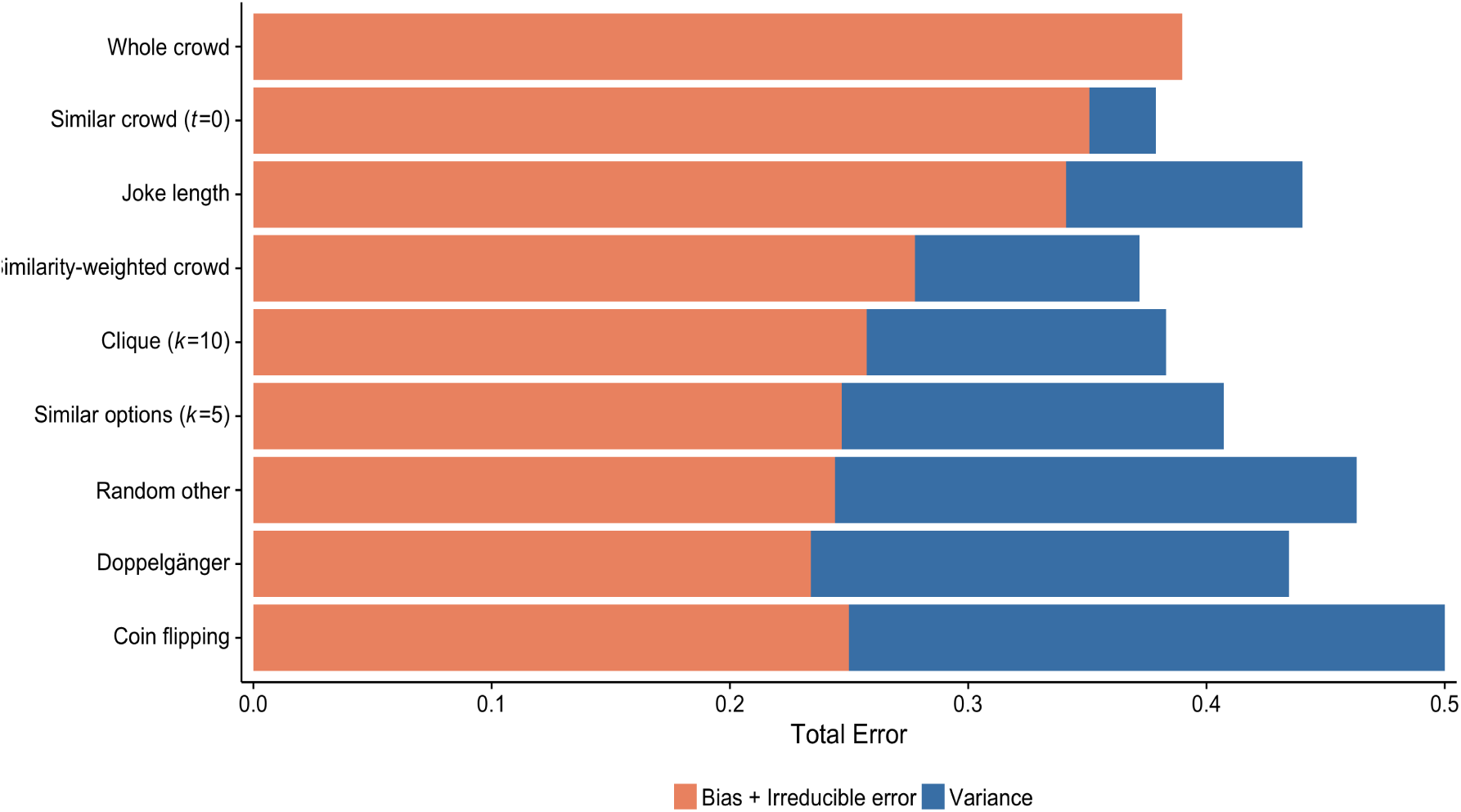
The bias–variance trade-off when we vary the numbers of people included in the clique. The figure shows the strategies’ performance with communities of 250 individuals when the models were trained on 25 jokes (based on the full matrix of evaluations). The bars represent the total misclassification error of the model (1 – % correct). The orange part of the bars represents the bias plus the irreducible error; the blue part represents the variance. In addition to the investigated strategies, we present the bias–variance decomposition of a coin-flipping algorithm as a measure of comparison.

We repeated the following procedure 500 times:

1. We divided the 14,000 individuals into 56 communities of 250 members each.
2. We divided the jokes into a training space *D* (75 jokes) and a test set *E* (25 jokes). This division was the same for all individuals in each community but differed across communities.
3. We generated all possible 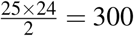 pair comparisons between jokes in the test set.
4. We randomly sampled uniformly without replacement 500 training sets of size *m* = 25 from the training space *D*. The training space was thus sufficiently bigger that the training sets *D* = 3*m*. Furthermore, because there are 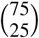 possible samples, the probability of sampling duplicate training sets was very small. We trained the models with each of these 500 samples.
5. The models estimated the funniness of each joke and chose the one with the higher estimate in each of the pair comparisons.
6. For each pair comparison, we calculated the variance component of the error. Subtracting variance from the total error, we calculated the *bias*^2^ + *irreducible error* component of the total error. For real-world data, the underlying data-generating distribution is rarely known. It is thus impossible to tease apart *bias*^2^ from *irreducible error*, and it is common practice to report them together [15]; see also our discussion of this point at the beginning of this section.

We then averaged the results over pair comparisons (300) and repetitions (100). In the following, we present average results, but we also break down the bias–variance profile of the investigated strategies for each individual in the population.

#### 10.8.6 Bias–Variance Analysis

##### Aggregate-level results

We start by discussing the results at the aggregate level. As shown in Figure S12, although the *whole crowd* strategy is the most biased of all the strategies, the variance component of the error is zero by design. As a result, the strategy has a much lower total error component than some of the less biased strategies (e.g., *similar options* and *doppelgänger*). Yet it is outperformed by the *similar crowd*, *similarity-weighted crowd*, and *clique* strategies, which are the best strategies at this level of experience. The *similar crowd* strategy is the second most biased strategy but has very little variance, leading to a relatively low error. The *similarity-weighted crowd* and *clique* strategies, which both rely on estimates of similarity to aggregate people’s evaluations, strike a good balance between bias and variance at this level of experience. In contrast, the error of the *doppelgänger* strategy, which relies on just one individual, has a particularly large variance component. The individual perceived as the most similar often changes across training samples, frequently leading to different predictions. Although the average predictions are close to the truth, they are also inconsistent, leading to a high variance component. Note that three strategies—*similar options*, *random other*, and *doppelgänger*—are less biased than coin flipping.^4^

##### The bias–variance trade-off within a single parametric strategy

Although the *doppelgänger*, the *clique*, and the *whole crowd* strategies are all special cases of the same recommender systems algorithm (k nearest neighbors; i.e., aggregating the opinions of the *k* most similar others), the previous analysis revealed substantial differences in their bias–variance profiles. At one extreme, the *whole crowd* strategy had the highest bias but no variance; at the other, the *doppelgänger* strategy had the lowest levels of bias, but very high variance (almost as high as flipping a coin at random). How does the bias–variance decomposition play out when we systematically vary the parameter *k* controlling the number of similar others whose opinions are aggregated? To investigate this question, we repeated the bias–variance simulation, systematically varying the number *k* of most similar others whose opinions are averaged. The simulation reveals a gradual decrease in bias as the number of people whose opinions are aggregated decreases. Yet this decrease is accompanied by a gradual increase in the variance of the strategy. The increase in variance is more pronounced when only very few similar others are considered (i.e., moving from the *clique* to the *doppelgänger* strategy). In this setting, the best bias–variance trade-off is achieved when the opinions of the 50 most similar others are aggregated.

**Figure S13:**
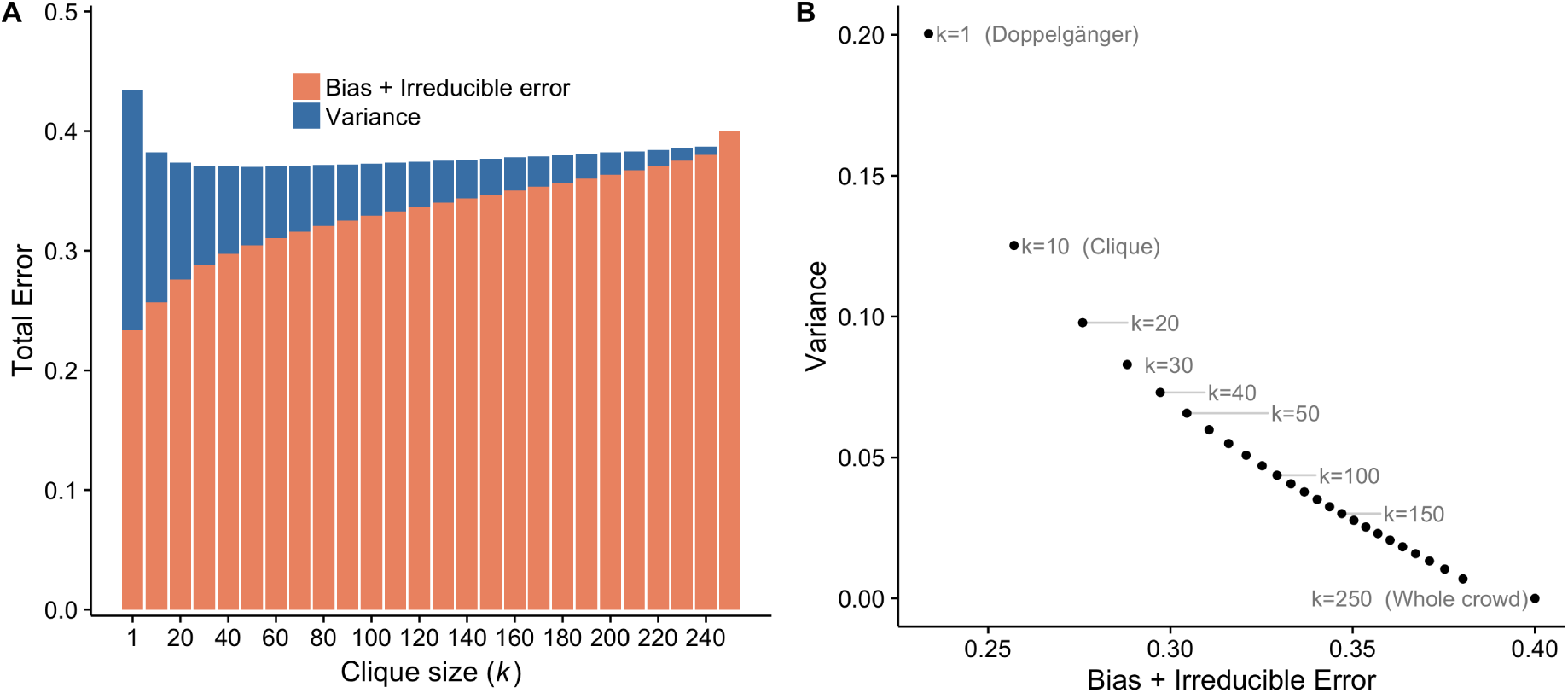
The bias–variance trade-off of social learning strategies. The figure shows the strategies’ performance with communities of 250 individuals when the models were trained on 25 jokes (based on the full matrix of evaluations). **Panel A:** The bars represent the total misclassification error (1 – proportion correct) of different instances of the clique model where we varied the number *k* of most similar others included in the clique. The orange part of the bars represents *bias*^2^+*irreducible error*; the blue part represents the variance. **Panel B:** A two-dimensional representation of the bias–variance decomposition when we vary the number *k* of most similar individuals included in the clique, where the x-axis shows *bias*^2^ +*irreducible error* and the y-axis shows variance. Recall that by setting *k* = 1 we get the *doppelgänger* strategy and by setting *k* = 250 we get the *whole crowd* strategy. The presented analysis decomposes the total error presented in Figure S8 for 25 previously experienced items.

##### Individual-level results

As discussed earlier, each individual learns his or her preferences in her own specific environment. As illustrated in Figure 2, performance varies markedly across individuals and depends both on their mean similarity with other people in the population and on the dispersion of similarities in their potential crowds (the standard deviation of the similarities). How does the total error decompose to bias and variance for different individuals? Figure S14 provides the answer to this question. For almost all of the strategies considered in this study, the variance component is almost the same across all individuals in the population (see the *doppelgänger*, *clique*, and *similar crowd* strategies). Thus, the differences in the bias of a strategy for different individuals show very similar patterns with those of the total error. All the strategies are more biased for individuals who have low mean similarity with the crowd and whose taste similarities with their peers are less dispersed. This effect is more striking for the aggregation strategies and marginally visible for the *doppelgänger* strategy. The *similarity-weighted crowd* strategy is the exception to this general pattern. For individuals with lower mean taste similarity, this strategy has a larger variance component in their error term. Furthermore, it has a larger bias for individuals whose peers have lower dispersion of taste similarity.

**Figure S14:**
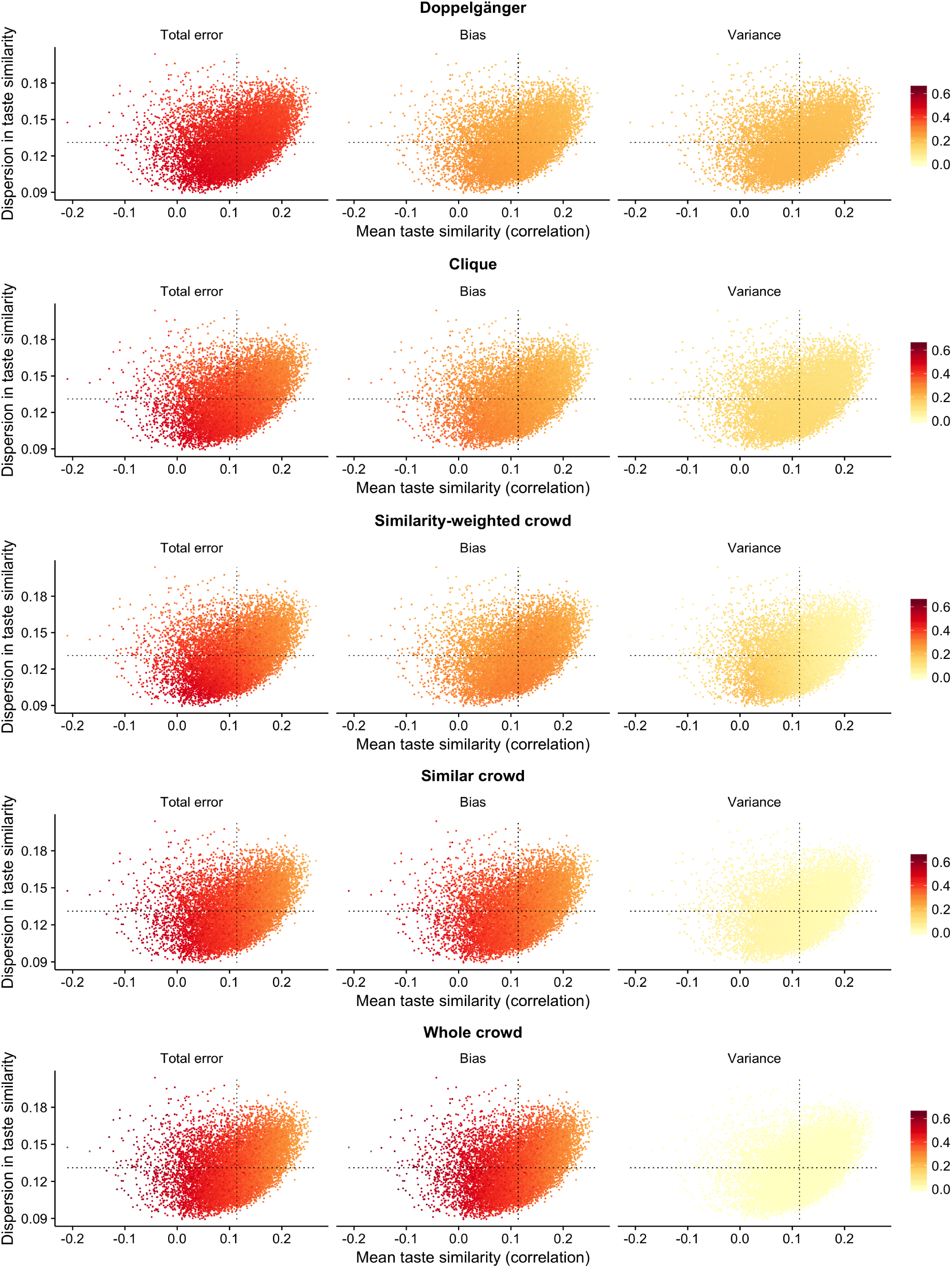
The bias–variance trade-off for five of the social learning strategies at the individual level. Each point represents one of the 14,000 individuals. Each individual is positioned according to their mean taste similarity with all other 13,999 individuals (x-axis) and the dispersion in taste similarity with other individuals (i.e., the standard deviation of these correlations; y-axis). The strategies are ordered from top to bottom according to their average variance at the aggregate level, starting from the *doppelgänger*, which has the highest variance, and closing with the *whole crowd*, which has no variance whatsoever.

## Footnotes

1 Some of the data are publicly available at http://files.grouplens.org/datasets/movielens/ml-10m-README.html

2 Although a coin toss is the paramount example of predicting at chance level in popular culture, it is in fact possible to access information that leads to better-than-chance predictions [2]. Later on we will break that assumption to enrich our illustrations.

3 When the number of items sampled in the training set is approximately the same as the entire pool of items, the estimates of variance are inevitably inaccurate.

4 Note that the Kohavi-Wolpert decomposition slightly overestimates the bias term of the error when relatively few training samples have been used to estimate the *bias* and *variance* components. This estimation error reduces to zero when one has the luxury to use many different training samples to calculate the components of the error. This was the case in our study. The authors also propose a method that can be used to reduce the estimation error when one lacks the computational capacity to repeat the simulation a large number of times.

## Reference

[1] Gediminas Adomavicius and Alexander Tuzhilin. Toward the next generation of recommender systems: A survey of the state-of-the-art and possible extensions. IEEE Transactions on Knowledge and Data Engineering, 17:734–749, 2005.

[2] Hirotogu Akaike. Information theory and an extension of the maximum likelihood principle. In Selected Papers of Hirotugu Akaike, pages 199–213. Springer, 1998.

[3] Naomi S Altman. An introduction to kernel and nearest-neighbor nonparametric regression. The American Statistician, 46(3):175–185, 1992.

[4] Pantelis P Analytis, Amit Kothiyal, and Konstantinos Katsikopoulos. Multi-attribute utility models as cognitive search engines. Judgment and Decision Making, 9(5):403–419, 2014.

[5] Sylvain Arlot, Alain Celisse, et al. A survey of cross-validation procedures for model selection. Statistics surveys, 4:40–79, 2010.

[6] Robert M Bell and Yehuda Koren. Lessons from the netflix prize challenge. Acm Sigkdd Explorations Newsletter, 9(2):75–79, 2007.

[7] Jesús Bobadilla, Fernando Ortega, Antonio Hernando, and Abraham Gutiérrez. Recommender systems survey. Knowledge-Based Systems, 46:109–132, 2013.

[8] Dirk Bollen, Bart P Knijnenburg, Martijn C Willemsen, and Mark Graus. Understanding choice overload in recommender systems. In Proceedings of the Fourth ACM Conference on Recommender Systems, pages 63–70. ACM, 2010.

[9] Robert Boyd and Peter J Richerson. Culture and the Evolutionary Process. University of Chicago Press, 1985.

[10] Leo Breiman. Arcing classifiers. Annals of Statistics, pages 801–824, 1998.

[11] Erica Briscoe and Jacob Feldman. Conceptual complexity and the bias/variance tradeoff. Cognition, 118(1):2–16, 2011.

[12] Timothy C Brock. Communicator-recipient similarity and decision change. Journal of Personality and Social Psychology, 1(6):650, 1965.

[13] Robin Burke. Hybrid recommender systems: Survey and experiments. User modeling and user-adapted interaction, 12(4):331–370, 2002.

[14] Robert E Burnkrant and Alain Cousineau. Informational and normative social influence in buyer behavior. Journal of Consumer Research, 2(3):206–215, 1975.

[15] Luigi Luca Cavalli-Sforza and Marcus W. Feldman. Cultural Transmission and Evolution: A Quantitative Approach. Princeton University Press, 1981.

[16] Robert T Clemen. Combining forecasts: A review and annotated bibliography. International Journal of Forecasting, 5(4):559–583, 1989.

[17] Joel B Cohen and Ellen Golden. Informational social influence and product evaluation. Journal of Applied Psychology, 56(1):54, 1972.

[18] Jason Dana and Robyn M Dawes. The superiority of simple alternatives to regression for social science predictions. Journal of Educational and Behavioral Statistics, 29(3):317–331, 2004.

[19] Harry L Davis, Stephen J Hoch, and EK Easton Ragsdale. An anchoring and adjustment model of spousal predictions. Journal of Consumer Research, 13(1):25–37, 1986.

[20] Clintin P Davis-Stober, David V Budescu, Jason Dana, and Stephen B Broomell. When is a crowd wise? Decision, 1(2):79–101, 2014.

[21] Clintin P Davis-Stober, Jason Dana, and David V Budescu. Why recognition is rational: Optimality results on single-variable decision rules. Judgment and Decision Making, 5(4):216, 2010.

[22] Robyn M Dawes. The robust beauty of improper linear models in decision making. American Psychologist, 34(7):571–582, 1979.

[23] Christian Desrosiers and George Karypis. A comprehensive survey of neighborhood-based recommendation methods. In Recommender Systems Handbook, pages 107–144. Springer, 2011.

[24] Robin Dunbar. How Many Friends Does One Person Need? Dunbar’s Number and Other Evolutionary Quirks. Faber & Faber, 2010.

[25] Casey M Eggleston, Timothy D Wilson, Minha Lee, and Daniel T Gilbert. Predicting what we will like: Asking a stranger can be as good as asking a friend. Organizational Behavior and Human Decision Processes, 128:1–10, 2015.

[26] Hillel J Einhorn and Robin M Hogarth. Unit weighting schemes for decision making. Organizational Behavior and Human Performance, 13(2):171–192, 1975.

[27] Hillel J Einhorn, Robin M Hogarth, and Eric Klempner. Quality of group judgment. Psychological Bulletin, 84(1):158–172, 1977.

[28] Michael D Ekstrand, F Maxwell Harper, Martijn C Willemsen, and Joseph A Konstan. User perception of differences in recommender algorithms. In Proceedings of the 8th ACM Conference on Recommender Systems, pages 161–168. ACM, 2014.

[29] Michael D Ekstrand, John T Riedl, and Joseph A Konstan. Collaborative filtering recommender systems. Foundation and Trends in Human-Computer Interaction, 4(2):81–173, 2011.

[30] Glenn Ellison and Drew Fudenberg. Rules of thumb for social learning. Journal of Political Economy, 101(4):612–643, 1993.

[31] Christine A Fawcett and Lori Markson. Children reason about shared preferences. Developmental Psychology, 46(2):299, 2010.

[32] Scott L Feld. Social structural determinants of similarity among associates. American sociological review, pages 797–801, 1982.

[33] Ignacio Fernández-Tobías, Matthias Braunhofer, Mehdi Elahi, Francesco Ricci, and Iván Cantador. Alleviating the new user problem in collaborative filtering by exploiting personality information. User Modeling and User-Adapted Interaction, 26(2-3):221–255, 2016.

[34] Jerome Friedman, Trevor Hastie, and Robert Tibshirani. The Elements of Statistical Learning: Data Mining, Inference, and Prediction. Springer, 2001.

[35] Stuart Geman, Elie Bienenstock, and René Doursat. Neural networks and the bias/variance dilemma. Neural Computation, 4(1):1–58, 1992.

[36] Samuel Gershman, Thomas Pouncy Hillard, and Hyowon Gweon. Learning the structure of social influence. Cognitive Science, 41(3), 2017.

[37] P Geurts. Bias vs variance decomposition for regression and classification. In Oded Maimon and Lior Rokach, editors, Data Mining and Knowledge Discovery Handbook, pages 733–746. Springer, 2010.

[38] Gerd Gigerenzer. From tools to theories: A heuristic of discovery in cognitive psychology. Psychological Review, 98(2):254–267, 1991.

[39] Gerd Gigerenzer and H Brighton. Homo heuristicus: Why biased minds make better inferences. Topics in Cognitive Science, 1(1):107–143, 2009.

[40] Gerd Gigerenzer and Daniel G Goldstein. Reasoning the fast and frugal way: Models of bounded rationality. Psychological Review, 103(4):650–669, 1996.

[41] Gerd Gigerenzer and Peter M Todd. Simple heuristics that make us smart. Oxford University Press, 1999.

[42] Daniel T Gilbert, Matthew A Killingsworth, Rebecca N Eyre, and Timothy D Wilson. The surprising power of neighborly advice. Science, 323(5921):1617–1619, 2009.

[43] Francesca Gino, Jen Shang, and Rachel Croson. The impact of information from similar or different advisors on judgment. Organizational Behavior and Human Decision Processes, 108(2):287–302, 2009.

[44] Ken Goldberg, Theresa Roeder, Dhruv Gupta, and Chris Perkins. Eigentaste: A constant time collaborative filtering algorithm. Information Retrieval, 4(2):133–151, 2001.

[45] Daniel G Goldstein, Randolph Preston McAfee, and Siddharth Suri. The wisdom of smaller, smarter crowds. In Proceedings of the Fifteenth ACM Conference on Economics and Computation, pages 471–488. ACM, 2014.

[46] Robert L Goldstone and Gary Lupyan. Discovering psychological principles by mining naturally occurring data sets. Topics in Cognitive Science, 8(3):548–568, 2016.

[47] Robert L Goldstone and Ji Yun Son. Similarity. Cambridge University Press, 2005.

[48] Robert L Goldstone, Thomas N Wisdom, Michael E Roberts, and Seth Frey. Learning along with others. Psychology of Learning and Motivation, 58:1–45, 2013.

[49] Kate Gordon. Group judgments in the field of lifted weights. Journal of Experimental Psychology, 7(5):398, 1924.

[50] Kenneth R Hammond, Carolyn J Hursch, and Frederick J Todd. Analyzing the components of clinical inference. Psychological Review, 71(6):438–456, 1964.

[51] Reid Hastie and Tatsuya Kameda. The robust beauty of majority rules in group decisions. Psychological Review, 112(2):494, 2005.

[52] Trevor Hastie, Robert Tibshirani, Jerome Friedman, and James Franklin. The elements of statistical learning: data mining, inference and prediction. The Mathematical Intelligencer, 27(2):83–85, 2005.

[53] Jonathan L Herlocker, Joseph A Konstan, Al Borchers, and John Riedl. An algorithmic framework for performing collaborative filtering. In Proceedings of the 22nd Annual International ACM SIGIR Conference on Research and Development in Information Retrieval, pages 230–237. ACM, 1999.

[54] Jonathan L Herlocker, Joseph A Konstan, Loren G Terveen, and John T Riedl. Evaluating collaborative filtering recommender systems. ACM Transactions of Information Systems, 22(1):5–53, 2004.

[55] Stefan M Herzog and Ralph Hertwig. The wisdom of many in one mind: Improving individual judgments with dialectical bootstrapping. Psychological Science, 20(2):231–237, 2009.

[56] Stefan M Herzog and Ralph Hertwig. Harnessing the wisdom of the inner crowd. Trends in Cognitive Science, 18(10):504–506, 2014.

[57] Stefan M Herzog and Bettina von Helversen. Strategy selection versus strategy blending: A predictive perspective on single- and multi-strategy accounts in multiple-cue estimation. Journal of Behavioral Decision Making.

[58] Robin M Hogarth. A note on aggregating opinions. Organizational Behavior and Human Performance, 21(1):40–46, 1978.

[59] Robin M Hogarth and Natalia Karelaia. Ignoring information in binary choice with continuous variables: When is less “more”? Journal of Mathematical Psychology, 49(2):115–124, 2005.

[60] David Hume. Of the standard of taste. 1757.

[61] Peter Juslin and Magnus Persson. PROBabilities from EXemplars (PROBEX): A “lazy” algorithm for probabilistic inference from generic knowledge. Cognitive Science, 26(5):563–607, 2002.

[62] Lee Jussim and D Wayne Osgood. Influence and similarity among friends: An integrative model applied to incarcerated adolescents. Social Psychology Quarterly, pages 98–112, 1989.

[63] Konstantinos V Katsikopoulos. Psychological heuristics for making inferences: Definition, performance, and the emerging theory and practice. Decision Analysis, 8(1):10–29, 2011.

[64] Vinod Krishnan, Pradeep Kumar Narayanashetty, Mukesh Nathan, Richard T Davies, and Joseph A Konstan. Who predicts better?: Results from an online study comparing humans and an online recommender system. In Proceedings of the 2008 ACM conference on Recommender systems, pages 211–218. ACM, 2008.

[65] John K Kruschke. ALCOVE: An exemplar-based connectionist model of category learning. Psychological Review, 99(1):22–44, 1992.

[66] Kevin N Laland. Social learning strategies. Animal Learning Behavior, 32(1):4–14, 2004.

[67] Richard P Larrick, Albert E Mannes, and Jack B Soll. The social psychology of the wisdom of crowds. In Joachim I Krueger, editor, Frontiers in Social Psychology: Social Judgment and Decision Making, pages 227–242. Psychology Press, New York, NY, 2012.

[68] Albert E Mannes, Jack B Soll, and Richard P Larrick. The wisdom of select crowds. Journal of Personality and Social Psychology, 107(2):276–299, 2014.

[69] Jutta Mata, Benjamin Scheibehenne, and Peter M Todd. Predicting children’s meal preferences: How much do parents know? Appetite, 50(2):367–375, 2008.

[70] Miller McPherson, Lynn Smith-Lovin, and James M Cook. Birds of a feather: Homophily in social networks. Annual review of sociology, 27(1):415–444, 2001.

[71] Barbara Mellers, Lyle Ungar, Jonathan Baron, Jaime Ramos, Burcu Gurcay, Katrina Fincher, Sydney E Scott, Don Moore, Pavel Atanasov, Samuel A Swift, et al. Psychological strategies for winning a geopolitical forecasting tournament. Psychological Science, 25(5):1106–1115, 2014.

[72] Johannes Müller-Trede, Shoham Choshen-Hillel, Meir Barneron, and Ilan Yaniv. The wisdom of crowds in matters of taste. Management Science, 2017.

[73] Robert M Nosofsky. Similarity scaling and cognitive process models. Annual Review of Psychology, 43(1):25–53, 1992.

[74] Maria Augusta SN Nunes and Rong Hu. Personality-based recommender systems: an overview. In Proceedings of the sixth ACM conference on Recommender systems, pages 5–6. ACM, 2012.

[75] Alexandra Paxton and Thomas L Griffiths. Finding the traces of behavioral and cognitive processes in big data and naturally occurring datasets. Behavioral Research Methods, pages 1–9, 2017.

[76] Michael Pazzani and Daniel Billsus. Content-based recommendation systems. In Peter Brusilovsky, Alfred Kobsa, and Wolfgang Nejdl, editors, The Adaptive Web, pages 325–341. Springer, 2007.

[77] Mark A Pitt and In Jae Myung. When a good fit can be bad. Trends in cognitive sciences, 6(10):421–425, 2002.

[78] Luke Rendell, Laurel Fogarty, William JE Hoppitt, Thomas JH Morgan, Mike M Webster, and Kevin N Laland. Cognitive culture: Theoretical and empirical insights into social learning strategies. Trends in Cognitive Science, 15(2):68–76, 2011.

[79] Betty M Repacholi and Alison Gopnik. Early reasoning about desires: Evidence from 14- and 18-month-olds. Developmental Psychology, 33(1):12, 1997.

[80] Paul Resnick, Neophytos Iacovou, Mitesh Suchak, Peter Bergstrom, and John Riedl. Grouplens: An open architecture for collaborative filtering of netnews. In Proceedings of the 1994 ACM Conference on Computer Supported Cooperative Work, pages 175–186. ACM, 1994.

[81] Paul Resnick and Hal R Varian. Recommender systems. Communications of the ACM, 40(3):56–58, 1997.

[82] Elaine Rich. User modeling via stereotypes. Cognitive Science, 3(4):329–354, 1979.

[83] Peter J Richerson and Robert Boyd. Not by Genes Alone: How Culture Transformed Human Evolution. University of Chicago Press, 2008.

[84] Jorma Rissanen. Modeling by shortest data description. Automatica, 14(5):465–471, 1978.

[85] Frank Rosenblatt. The perceptron: A probabilistic model for information storage and organization in the brain. Psychological Review, 65(6):386, 1958.

[86] Badrul Sarwar, George Karypis, Joseph Konstan, and John Riedl. Application of dimensionality reduction in recommender system-a case study. Technical report, Minnesota Univ Minneapolis Dept of Computer Science, 2000.

[87] Badrul Sarwar, George Karypis, Joseph Konstan, and John Riedl. Item-based collaborative filtering recommendation algorithms. In Proceedings of the 10th International Conference on World Wide Web, pages 285–295. ACM, 2001.

[88] Tobias Schnabel, Adith Swaminathan, Ashudeep Singh, Navin Chandak, and Thorsten Joachims. Recommendations as treatments: debiasing learning and evaluation. In Proceedings of the 33rd International Conference on International Conference on Machine Learning-Volume 48, pages 1670–1679. JMLR. org, 2016.

[89] Gideon Schwarz et al. Estimating the dimension of a model. The annals of statistics, 6(2):461–464, 1978.

[90] Upendra Shardanand and Pattie Maes. Social information filtering: Algorithms for automating “word of mouth”. In Proceedings of the SIGCHI Conference on Human Factors in Computing Systems, pages 210–217. ACM Press/Addison-Wesley Publishing Co., 1995.

[91] Amit Sharma and Dan Cosley. Studying and modeling the connection between people’s preferences and content sharing. In Proceedings of the 18th ACM Conference on Computer Supported Cooperative Work & Social Computing, pages 1246–1257. ACM, 2015.

[92] Herbert W Simons, Nancy N Berkowitz, and R John Moyer. Similarity, credibility, and attitude change: A review and a theory. Psychological Bulletin, 73(1):1–16, 1970.

[93] Özgür Şimşek. Linear decision rule as aspiration for simple decision heuristics. In Advances in Neural Information Processing Systems, pages 2904–2912, 2013.

[94] Özgür Şimşek and Marcus Buckmann. Learning from small samples: An analysis of simple decision heuristics. In Advances in Neural Information Processing Systems, pages 3141–3149, 2015.

[95] Rashmi R Sinha and Kirsten Swearingen. Comparing recommendations made by online systems and friends. In DELOS Workshop: Personalisation and Recommender Systems in Digital Libraries, volume 106, 2001.

[96] Louis L Thurstone. A law of comparative judgment. Psychological Review, 34(4):273–286, 1927.

[97] Jack L Treynor. Market efficiency and the bean jar experiment. Financial Analysis Journal, 43(3):50–53, 1987.

[98] Vladimir Naumovich Vapnik and Vlamimir Vapnik. Statistical learning theory, volume 1. Wiley New York, 1998.

[99] Patricia M West. Predicting preferences: An examination of agent learning. Journal of Consumer Research, 23(1):68–80, 1996.

[100] Robert L Winkler and Spyros Makridakis. The combination of forecasts. Journal of the Royal Statistical Society A, pages 150–157, 1983.

[101] Ilan Yaniv, Shoham Choshen-Hillel, and Maxim Milyavsky. Receiving advice on matters of taste: Similarity, majority influence, and taste discrimination. Organizational Behavior and Human Decision Processes, 115(1):111–120, 2011.

[102] Michael Yeomans, Anuj K Shah, Sendhil Mullainathan, and Jon Kleinberg. Making sense of recommendations.

[103] Philip W Yetton and Preston C Bottger. Individual versus group problem solving: An empirical test of a best-member strategy. Organizational Behavior and Human Performance, 29(3):307–321, 1982.

## Reference

[1] Leo Breiman. Bias, variance, and arcing classifiers. Technical report, 1996.

[2] Persi Diaconis, Susan Holmes, and Richard Montgomery. Dynamical bias in the coin toss. SIAM Review, 49(2):211–235, 2007.

[3] Robin Dunbar. How Many Friends Does One Person Need? Dunbar’s Number and Other Evolutionary Quirks. Faber & Faber, 2010.

[4] Michael D Ekstrand, John T Riedl, and Joseph A Konstan. Collaborative filtering recommender systems. Foundation and Trends in Human-Computer Interaction, 4(2):81–173, 2011.

[5] Scott Fortmann-Roe. Understanding the bias-variance tradeoff. http://scott.fortmann-roe.com/docs/BiasVariance.html. Accessed: 2017-07-29.

[6] Jerome H Friedman. On bias, variance, 0/1—loss, and the curse-of-dimensionality. Data Mining and Knowledge Discovery, 1(1):55–77, 1997.

[7] Stuart Geman, Elie Bienenstock, and René Doursat. Neural networks and the bias/variance dilemma. Neural Computation, 4(1):1–58, 1992.

[8] Arthur E Hoerl and Robert W Kennard. Ridge regression: Biased estimation for nonorthogonal problems. Technometrics, 12(1):55–67, 1970.

[9] Robin M Hogarth and Natalia Karelaia. Ignoring information in binary choice with continuous variables: When is less “more”? Journal of Mathematical Psychology, 49(2):115–124, 2005.

[10] Gareth M James. Variance and bias for general loss functions. Machine Learning, 51(2):115–135, 2003.

[11] Ron Kohavi and David H Wolpert. Bias plus variance decomposition for zero-one loss functions. In Proceedings of the Thirteenth International Conference on Machine Learning, pages 275–83, 1996.

[12] Eun Bae Kong and Thomas G Dietterich. Error-correcting output coding corrects bias and variance. In Proceedings of the Twelfth International Conference on Machine Learning, pages 313–321, 1995.

[13] Özgür Şimşek and Marcus Buckmann. On decision heuristics, bias, and variance. Manuscript in preparation, 2016.

[14] Robert Tibshirani. Bias, variance and prediction error for classification rules, 1996.

[15] Peter Van Der Putten and Maarten Van Someren. A bias-variance analysis of a real world learning problem: The CoIL Challenge 2000. Machine Learning, 57(1-2):177–195, 2004.

